# Juvenile hormones direct primordial germ cell migration to the embryonic gonad

**DOI:** 10.1101/2021.09.30.462471

**Authors:** Barton Lacy J, Sanny Justina, Dawson Emily P, Nouzova Marcela, Noriega Fernando Gabriel, Stadtfeld Matthias, Lehmann Ruth

**Affiliations:** Department of Cell Biology, Skirball Institute of Biomolecular Medicine, and Howard Hughes Medical Institute, NYU Grossman School of Medicine, NY, USA, 540 First Avenue, New York, NY 10016, USA; Department of Neuroscience, Developmental and Regenerative Biology, The University of Texas at San Antonio, One UTSA Circle, San Antonio, TX, 78249, USA; Department of Biological Sciences and Biomolecular Sciences Institute, Florida International University, 11200 SW 8^th^ St, Miami, FL 33199, USA; Institute of Parasitology, Biology Centre CAS, Ceske Budejovice, Czech Republic; Department of Parasitology, University of South Bohemia, České Budějovice, 37005, Czech Republic; Present: Department of Medicine, Weill Cornell Medicine, 413 E 69^th^ St, New York, NY 10021, USA; Present: Whitehead Institute and Department of Biology, MIT, 455 Main Street, Cambridge, MA 02142, USA

**Keywords:** primordial germ cells, cell migration, gonad, germline, mevalonate pathway, juvenile hormone, retinoids

## Abstract

Germ cells are essential to sexual reproduction. Across the animal kingdom, extracellular signaling isoprenoids, such as retinoic acids (RAs) in vertebrates and juvenile hormones (JHs) in invertebrates, facilitate multiple processes in the germline lifecycle. Here we investigated the role of these potent signaling molecules in embryonic germ cell migration, using JHs in *Drosophila melanogaster* as a model system. In contrast to their established endocrine roles during larval and adult germline development, we found that JH signaling acts locally during embryonic development. Using an *in vivo* biosensor, we found JH signaling is first active near primordial germ cells (PGCs) as they migrate to the developing somatic gonad. Through *in vivo* and *in vitro* assays, we found that JHs are both necessary and sufficient for PGC migration. Analysis into the mechanisms of this newly uncovered paracrine JH function revealed that PGC migration was compromised when JHs were reduced or increased, suggesting that specific titers or spatiotemporal JH dynamics are required for robust PGC colonization to the gonad. Compromised PGC migration can impair fertility and cause germ cell tumors in many species, including humans. In mammals, retinoids, a JH-related family of signaling isoprenoids, has many roles in development and reproduction. We found that retinoic acid, like JH, was sufficient to impact PGC migration *in vitro*. Together, our study reveals a previously unanticipated role of isoprenoids as local effectors of pre-gonadal PGC development and suggests a broadly shared mechanism in PGC migration.

**Highlights:**

- Juvenile hormones (JH) are necessary & sufficient for Primordial Germ Cell (PGC) migration.
- JH signaling acts directly in and around migrating PGCs prior to its endocrine function.
- Compensatory feedback sensitive to JH receptor function maintains JH homeostasis.
- JH-like retinoic acids may have similar roles during mammalian germ cell migration.

## Introduction

In sexually reproducing species, the germline links one generation to the next. Given the importance to individuals, societies, and species, understanding how germ cells are set aside, develop, and protected until undergoing gametogenesis to yield sperm and egg has been the focus of intense efforts. Through recent advances in single-cell ‘omics and *in vitro* derivation of primordial germ cell (PGC)-like cells from induced pluripotent stem cells, we have gained substantial insights into the molecular programs intrinsically required for germ cell specification and development. However, the yield and gamete quality of *in vitro* derivation approaches falls far short of what is observed *in vivo*^1^. This quality gap suggests that we are failing to recapitulate key aspects of germline development. One critical component not yet replicated *in vitro*, nor fully understood *in vivo*, is how germline development is exquisitely coordinated in space and time during embryogenesis by surrounding somatic cells.

Somatic cells coordinate germ cell development and gametogenesis through small, secreted molecules such as steroid hormones and signaling lipids. These potent molecules have multifaceted impacts on reproduction across the animal kingdom. In vertebrate animals, key reproductive small molecules include steroid hormones (i.e., estrogen and testosterone) that are synthesized from cholesterol, and paracrine retinoic acids (RAs), that are derived from carotenoids, a subfamily of isoprenoids. Animals lack the ability to synthesize carotenoids and thus obtain these critical molecules from their diet. Despite differences in biosynthetic origins and structures, invertebrate animals also use steroids and isoprenoids to facilitate many aspects of germline development and reproduction. In *Drosophila*, the steroid hormone, ecdysone, regulates germline stem cell homeostasis, niche development, yolk uptake, and the migratory behavior of certain ovarian somatic cell types^2–5^. The RA-like isoprenoids, Juvenile hormones, have profound impacts on reproduction. For example, methyl farnesoate is a key sex determination molecule in some crustaceans^6^. In many insects, JHs act as gonadotropins, with essential roles in vitellogenesis^7, 8^. Recent work in *Drosophila* have revealed that JHs also regulate germline stem cell number and reproductive diapause^9, 10^.

While much is known about how steroid hormones mechanistically shepherd the germline lifecycle, how signaling isoprenoids regulate the germline lifecycle is much less understood. Given the complexity surrounding RAs in germline biology, insights can be gleaned from JHs. These potent sesquiterpenoids, derived from the mevalonate pathway^11^, are best known for their influence on juvenile development^12^. They include, among others, JH III, JH III bisepoxide (JHB_3_), and methyl farnesoate (**Fig 1a**). As signaling molecules, JHs bind to a basic helix-loop helix/Per-ARNT-SIM (bHLH-PAS) receptor, which in *Drosophila* is encoded by two paralogous genes, *Methoprene-tolerant* (*Met*) and *germ cell-expressed bHLH-PAS* (*gce*)^13–15^. Upon ligand binding, these receptors regulate gene expression together with the co-activator Taiman^16^. In contrast to paracrine acting RAs, JHs act hormonally such that JH is synthesized in the corpus allatum (CA), a neuroendocrine gland, and released into the circulatory system (**Fig 1a**). JH production is driven by Juvenile hormone acid methyltransferase (Jhamt), which is expressed in the CA as soon as this gland forms at the end of embryogenesis^17^. Curiously, a burst of *jhamt* expression has also been observed in the trunk mesoderm during mid-embryogenesis, well before the specification of the CA precursors^17^, leaving open the possibility that JHs have additional, non-endocrine roles.

**Figure 1:**
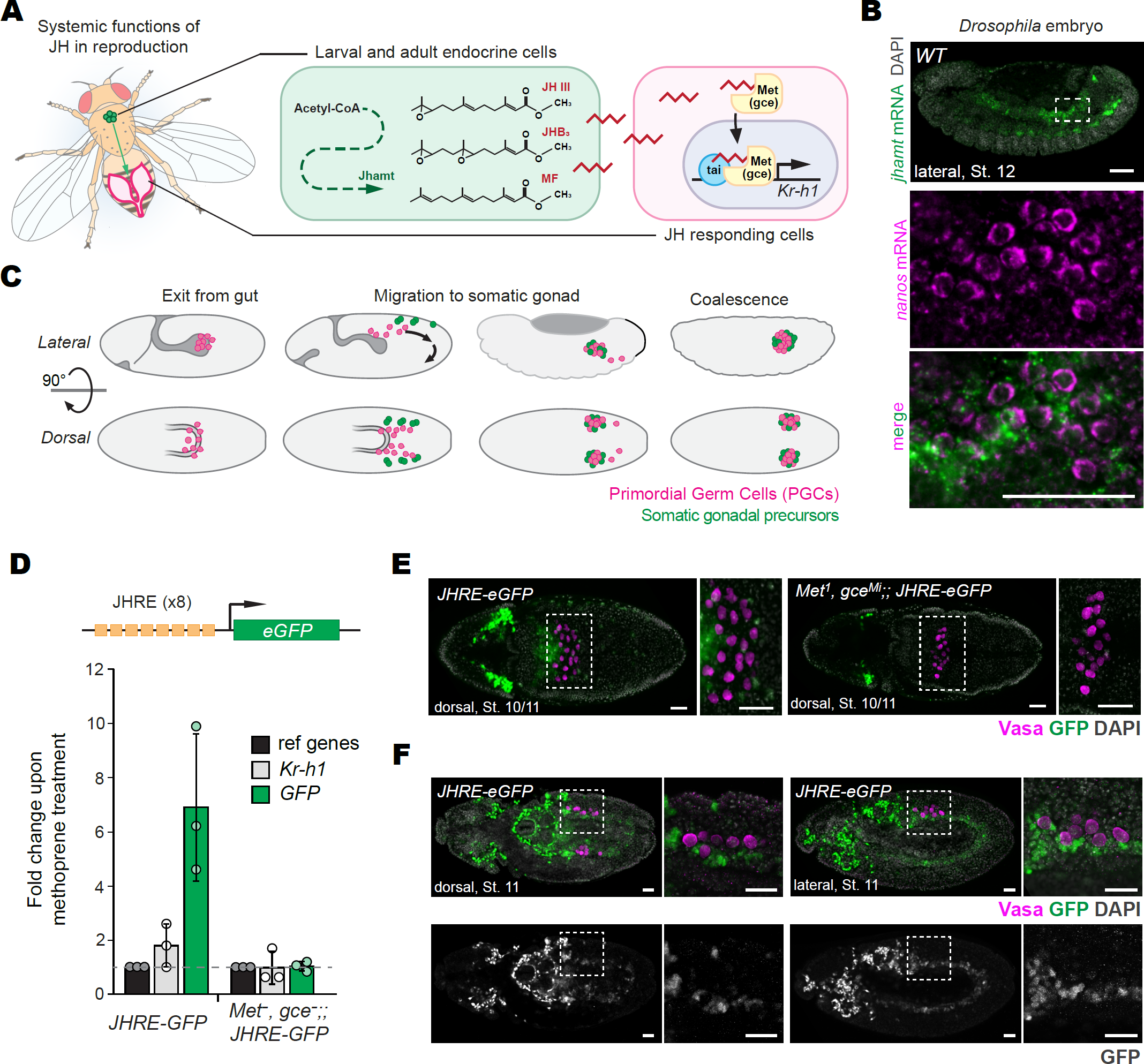
Juvenile hormone (JH) signaling is active in the embryonic mesoderm during PGC migration. **A** *Left*: Schematic summarizing gonadotropic effect of JH in insects, whereby JH is produced in an endocrine gland and acts systemically to promote oogenesis. *Right*: Summary of established JH signaling pathways wherein endocrine cells produce JHs downstream of Acetyl-CoA and a biosynthetic pathway that culminates in JH acid methyltransferase (Jhamt)-mediated JH production. JHs (red jagged lines) are then secreted in the hemolymph and enter JH-responding cells. Once in JH-responding cells, JH binds to one of two JH transcriptional receptors, Methoprene tolerant (Met) or Germ cell expressed (Gce). JH-bound Met/Gce translocates to the nucleus to regulate target genes. **B** Lateral view images of a late Stage 12 embryo stained by Fluorescent *in situ* hybridization (FISH) for *jhamt* (green) and *nanos* (magenta) mRNA. Primers used to generate *in situ* probes are listed in Supplementary Table 3. *Right:* Magnified image of *jhamt* and *nanos* mRNA in the boxed area outlined by a dashed, white square. **C** Schematic of germ cell development during Stage 10-14 of *Drosophila* embryogenesis. **D** *Top*: schematic of a new transgenic JH sensor (JHRE-GFP) in which eight copies of the JH Response Element from the *early trypsin* gene from *A. aegypti* were placed upstream of eGFP (See Materials and Methods for construct design). *Bottom*: Induction of *Kr-h1* mRNA, a known JH transcriptional target, and *GFP* mRNA. deltaCt was calculated relative to the average Ct of two housekeeping genes, *DCTN5-p25* and *Und*, and fold change was calculated for whole second instar larvae fed cornmeal food containing 1.6µg/mg of JH analogue, methoprene normalized to age and genotype-matched larvae fed ethanol. Animals were fed methoprene or solvent control for 24 hours prior to RNA isolation. Bars and error bars represent average fold change and standard deviation from three biological replicates, each consisting of ten second instar larvae. **E** Dorsal (d) images of Stage 10 *JHRE-GFP* embryos that are wild type (left) or mutant for Met and Gce (right). **F** *Left:* dorsal view of a Stage 11 *JHRE-GFP* embryo stained for GFP (green) protein, Vasa (magenta) protein, and DNA using DAPI (gray). *Right*: lateral view of a Stage 11 JHRE-GFP embryo. GFP channel is shown below each merged image. For all images, magnified images in the boxed area outlined by a dashed, white square are shown to the right of each whole-embryo image. Genotypes are noted in the top left corner, embryo orientation in the bottom, and scale bars represent 25µm. All images are maximum projections of two to three 2µm confocal slices.

Mesodermal *jhamt* expression coincides with the PGC migration stage of reproductive development. In many species, newly specified PGCs are internalized during gastrulation and then must migrate within the mesoderm toward the developing gonad to generate the germline pool. In some mammals, including humans, extragonadal PGCs that fail to reach the gonad can cause germ cell tumors^18^. To investigate the role of JHs during PGC migration, we first generated an *in vivo* JH sensor and found that JH signaling was first active proximal to *jhamt* expression and surrounding migratory PGCs. New *jhamt* mutants lacking the methyltransferase domain compromised JH synthesis and caused a failure of PGCs reaching the gonad. A similar defect in gonad colonization had previously been described in animals lacking HMG Coenzyme A reductase (Hmgcr)^19^, the rate-limiting enzyme of the mevalonate pathway upstream of Jhamt in JH synthesis^11^. Consistent with Hmgcr and Jhamt being part of the same synthetic cascade, we found that *jhamt* and *Hmgcr* genetically interact to facilitate PGC migration and that JH, like Hmgcr, is both necessary and sufficient for PGC migration. Our further investigations into the function of JH receptors in the embryo revealed a feedback mechanism necessary to maintain JH homeostasis critical for PGC migration. Together, our studies uncovered a new requirement for JH signaling in embryonic reproductive development. Studies in mice demonstrated that, like JHs, genes critical to RA synthesis are expressed in the bipotential gonad during PGC development PGC development^20–22^. However, despite shared expression of genes involved in reproductive isoprenoids in vertebrate and invertebrate species, a role for isoprenoid signaling during the PGC migration has not been described. Here we demonstrate that RA is sufficient to induce migration of both *Drosophila* and mouse embryonic germ cells, suggesting this newly identified role for secreted isoprenoids on early germline development may be broadly shared across invertebrate and vertebrate species.

## Results

### Juvenile hormone signaling is active in the embryonic mesoderm during PGC development

Juvenile hormone production is driven by Juvenile hormone acid methyltransferase (Jhamt). Jhamt is specifically expressed in the corpus allatum (CA), as soon as this JH-producing endocrine gland develops in late embryogenesis^17^. Curiously, in *Drosophila jhamt* mRNA is also detected during mid-embryogenesis in the trunk mesoderm, before specification and migration of CA precursors^17^. Using dual-label fluorescent *in situ* hybridization for mRNAs encoding Jhamt and the germ cell factor, Nanos, we investigated where *jhamt* is expressed relative to PGCs. Consistent with previous work^17^, we observed *jhamt* expression in the mesoderm from embryonic Stage 10 (∼4 hours after fertilization), continuing through Stage 13 (∼11 hours after fertilization) (**Fig 1b**). At this stage in embryogenesis, PGCs exit the endoderm and sort into two lateral populations as they migrate within the mesoderm toward the somatic gonadal precursors (**Fig 1c**). This *jhamt* expression pattern suggested that JH synthesis may occur prior to CA formation and be involved in embryonic germline development.

To determine whether mesodermal *jhamt* mRNA leads to functional JH production during *Drosophila* embryogenesis, we generated an *in vivo* JH sensor. We placed *eGFP* downstream of eight tandem copies of a JH response element (JHRE) from the *early trypsin* gene of *Aedes aegypti*^15, 23^, which was previously shown to robustly and specifically responds to JH *in vitro*^15^ (**Fig 1d and Supp Fig 1a**). To test the responsiveness of this biosensor to JH *in vivo*, we fed second instar larvae the JH mimic, methoprene, and measured mRNA levels of *GFP* and the canonical JH signaling transcriptional target, *Krüppel homolog 1* (*Kr-h1*), by real-time-qPCR 24 hours later. We found that *Kr-h1* and *GFP* mRNA levels increased ∼two-fold and seven-fold respectively in *JHRE-GFP* larvae fed methoprene (**Fig 1d**). To determine whether this induction by methoprene required JH signaling, we generated animals carrying the JHRE sensor but lacking the JH receptors, Methoprene-tolerant (Met) and Germ cell-expressed bHLH-PAS (gce). We found that methoprene did not increase expression of either *Kr-h1* or *GFP* mRNA in *Met, gce; JHRE-eGFP* mutant animals (**Fig 1d**). Together, these data indicate that this new JH sensor faithfully responds to JH signaling *in vivo*.

With this *in vivo* JH sensor, we next investigated if and where JH signaling is active during embryonic germline development. We stained *JHRE-GFP* embryos with antibodies against GFP and the germ cell-specific RNA helicase, Vasa. GFP signal was first detected in Stage 10 embryos in two locations: in bilateral clusters in the anterior portion of the embryo and in bilateral strips of the mesoderm of the posterior germband (**Fig 1e, f**). The anterior signal was still detected in *Met, gce* mutant embryos carrying the JH biosensor, but the germband signal was absent (**Fig 1e**). To explore whether this difference is due to non-specific, receptor-independent transgene activation, we analyzed animals carrying a JH sensor transgene in which the Met/gce binding motif was mutated (**Supp Fig 1a**)^15^. Anterior GFP was still present in animals with the mutated biosensor; further investigation revealed that this non-JH-specific GFP signal localized to hemocytes (**Supp Fig 1b-c**). Importantly, the mesodermal signal was not detected in embryos carrying the JH biosensor with mutated receptor binding motifs, nor when the functional biosensor JHRE-eGFP was tested in *Met, gce* receptor mutants. Thus, the mesodermal signal is due to JH-mediated receptor activation and reflects JH signaling. We found that this specific JH sensitive response began at Stage 10 in the mesoderm, when PGCs enter this tissue from the endoderm. JH signaling continued to localize in and near PGCs as they colonized the gonad (**Fig 1e, Supp 1c**). Together, these data demonstrate that JH signaling is active near the mesodermal *jhamt* expression zone and surrounds migratory PGCs during mid-embryogenesis.

### Juvenile hormones are required for migrating PGCs to reach the somatic gonad

To determine whether the newly identified embryonic JH signaling supports early PGC development, we generated new mutant *jhamt* alleles. Using CRISPR editing, 15 *jhamt* alleles were generated, two of which carried a premature stop codon at amino acid 46, whereas the remaining 13 carried a premature stop codon at amino acid 54 (**Supp Fig 2**). All new alleles caused a deletion of the methyl transferase domain central to Jhamt-mediated JH biosynthesis (**Fig 2a**). To avoid confounding effects from second site mutations, phenotypic analyses carried out in this study use the newly generated *jhamt* lines in heteroallelic combinations or *in trans* to an independently generated small deficiency lacking the entire *jhamt* locus (**Supp Table 1**). To verify that Jhamt function was compromised, titers of two JHs in *Drosophila*, JH III and JHB_3_, were directly quantified by liquid chromatography-tandem mass spectrometry^24^. We found that JH III titers were reduced 50% and JHB_3_ titers were reduced >90% in larvae possessing our newly generated alleles (**Fig 2b**). These results are consistent with previous observations showing that JHs were still detected in animals lacking the entire *jhamt* locus^25^, suggesting the *Drosophila* genome may contain a second, yet obscure, methyltransferase capable of JH synthesis^26, 27^. To attest the degree by which JH function was compromised, we measured female fecundity. Consistent with previous reports^25^, the number of eggs laid per day was significantly reduced in females carrying our new *jhamt* mutant alleles (**Fig 2c**). Together, these data suggest that the newly generated *jhamt* mutants significantly affected JH production and display JH loss of function phenotypes.

**Figure 2:**
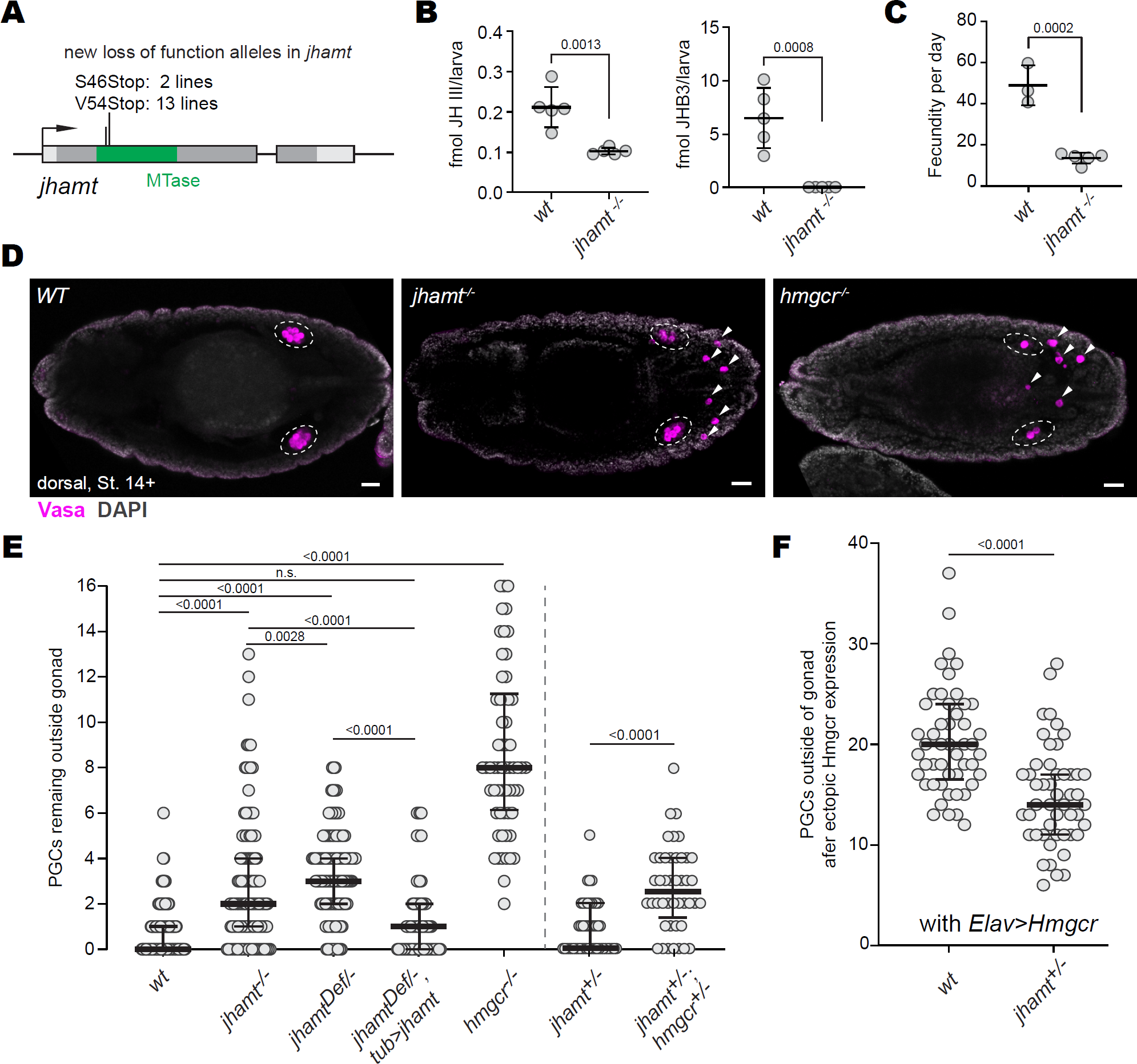
Key JH synthesis enzyme, Jhamt, facilitates PGC migration to the gonad. **A** Schematic of the *jhamt* locus noting the location of premature stop codons within the methyltransferase domain (shown in green), which were introduced by CRISPR-mediated editing. Light gray denotes 5’ and 3’ UTR. **B** Titers of two JHs, JH III and JH bisepoxide (JHB_3_), in hemolymph of third instar *Drosophila* larvae measured by quantitative mass spectrometry using deuterated standards. Each dot represents a biological replicate of hemolymph bled from 25 larvae. For titers, *w^−^, Sp/CyO,ft-lacZ* was used as a wildtype strain as the CRISPR-induced alleles in a w^−^ background were initially crossed into this background. **C** Fecundity of young females. Each dot represents the average number of eggs laid per biological replicate, which was obtained by dividing the number of eggs laid over a 24-hour period by the number of females (10-12) in each cohort three- and four-days after eclosion. To avoid confounding male infertility effects, all virgin females were mated to wildtype (*w*^1118^) males. For B and C, bars and whiskers represent mean +/-standard deviation and statistical significance was tested by unpaired student t test. **D** Dorsal view images of Stage 14 *Drosophila* embryos stained for Vasa (magenta) and DNA (DAPI, gray). Zygotic genotypes are noted in the top left corner. Gonads are circled with a white dashed circle. Arrowheads highlight extragonadal germ cells. All embryo images are maximum projections of two to three 2µm confocal slices. Scale bars represent 25µm. **E** Quantification of extragonadal PGCs in Stage 14-16 embryos. Each dot represents one embryo. To yield *jhamt^−/−^* embryos, with heteroallelic combinations of independently derived indels, *jhamt^8F^*^1^*/CyO,ftz-lacZ* females were mated to *jhamt^12F^*^1^*/CyO,ftz-lacZ*. To generate *jhamt^−/Def^* embryos, *jhamt^8F^*^1^*/CyO,ftz-lacZ* animals were mated to *Df(2L)BSC781/CyO, ft-lacZ* animals. **F** Quantification of extragonadal PGCs in Stage 14-16 embryos when Hmgcr is ectopically expressed in the nervous system using *Elav-GAL4* and *UAS-Hmgcr*. Each dot represents an embryo. For panels E and F, bars and whiskers represent median +/-interquartile range (IQR), statistical significance was tested by Mann-Whitney U test, with ns=not significant. The reference strain for *jhamt* studies was *w*^1118^ as *jhamt* alleles were backcrossed into this background.

With new genetic tools, we next compared PGC development in wildtype and *jhamt* mutant embryos. The first detectable germline defect in *jhamt* mutant embryos was an increased number of PGCs that failed to reach the gonad at Stage 14, when PGC normally have completed their migration to the somatic gonad (**Fig 2d)**. Animals lacking zygotic Jhamt had significantly more extragonadal PGCs than wildtype animals (**Fig 2e**). This phenotype was rescued when *jhamt* was zygotically expressed with a broad *Tub-GAL4* driver (**Fig 2e**). These data indicate that *jhamt* expressed in somatic cells facilitates PGC migration to the developing somatic gonad.

The *jhamt* mutant PGC phenotype was similar to that of embryos lacking HMG-CoA reductase (Hmgcr), the rate-limiting enzyme in the mevalonate pathway (**Fig 2d** and **e**)^19^. In *Drosophila*, Hmgcr was shown to have a striking, instructive effect on PGC migration: in *Hmgcr* mutant embryos PGCs failed to reach the gonad and when mis-expressed, elevated *Hmgcr* expression alone mislocalized PGCs to the location of ectopic expression (**Fig 2f** and **Supp Fig 2a, b)**^19, 28, 29^. Our observation that *jhamt* mutants phenocopied *Hmgcr* mutants is consistent with previous reports that Hmgcr is required for JH synthesis and genetically interacts with *jhamt* in the context of animal development^25^. To determine whether Jhamt functions downstream of Hmgcr in embryonic PGC migration, we tested for genetic interaction in two ways. First, we measured the effect of genetic *Hmgcr* reduction in sensitized *jhamt* heterozygous animals and found that significantly more PGCs failed to reach the gonad in *jhamt^+/-^; Hmgcr^+/-^*heterozygous embryos than embryos heterozygous for *jhamt* alone (**Fig 2e**). Second, we tested whether genetic reduction of Jhamt can suppress the ability of ectopic *Hmgcr* expression to misdirect PGCs. Indeed, loss of one wildtype *jhamt* copy significantly reduced the number of PGCs misdirected by GAL4/UAS-mediated expression of *Hmgcr* (**Fig 2f**). Together, these data suggest that Jhamt functions downstream of Hmgcr to facilitate PGC migration to the somatic gonad.

### Juvenile hormone is sufficient for PGC migration

Hmgcr is sufficient to misdirect migrating PGC to many tissues, even tissues far from the normal migratory path such as the central nervous system (**Fig 3a** and **Supp Fig 3a**)^19, 28, 29^. To determine whether, like *Hmgcr, jhamt* expression is sufficient for PGC migration *in vivo*, we ectopically expressed *jhamt* mRNA using the GAL4/UAS system. We found that neuronal *Elav-GAL4* driven *jhamt* expression led to modest, yet significant, increase in PGCs inappropriately near neuronal Elav-positive cells (**Fig 3b**). We even found some PGCs localized in the central nervous system after neuronal *jhamt* expression (**Fig 3a**). Yet overall, significantly fewer PGCs were misdirected by ectopic *jhamt* compared to ectopic *Hmgcr* driven by the same *Elav-GAL4* transgene (**Fig 3b**). Moreover, ectopic *jhamt* expression was insufficient to drive PGCs to inappropriate locations when *UAS-jhamt* was induced by the endodermal driver, *48Y-GAL4*, or the pan-mesodermal drivers, *twist-GAL4* and *24B-GAL4*, even though ectopic *Hmgcr* expression with these GAL4 drivers significantly mislocalized PGCs (**Supp Fig 3b**). These data show that Jhamt was sufficient to impact PGC migration *in vivo* in at least one, but not all, tissues, suggesting that other pathway components may be limiting.

**Figure 3:**
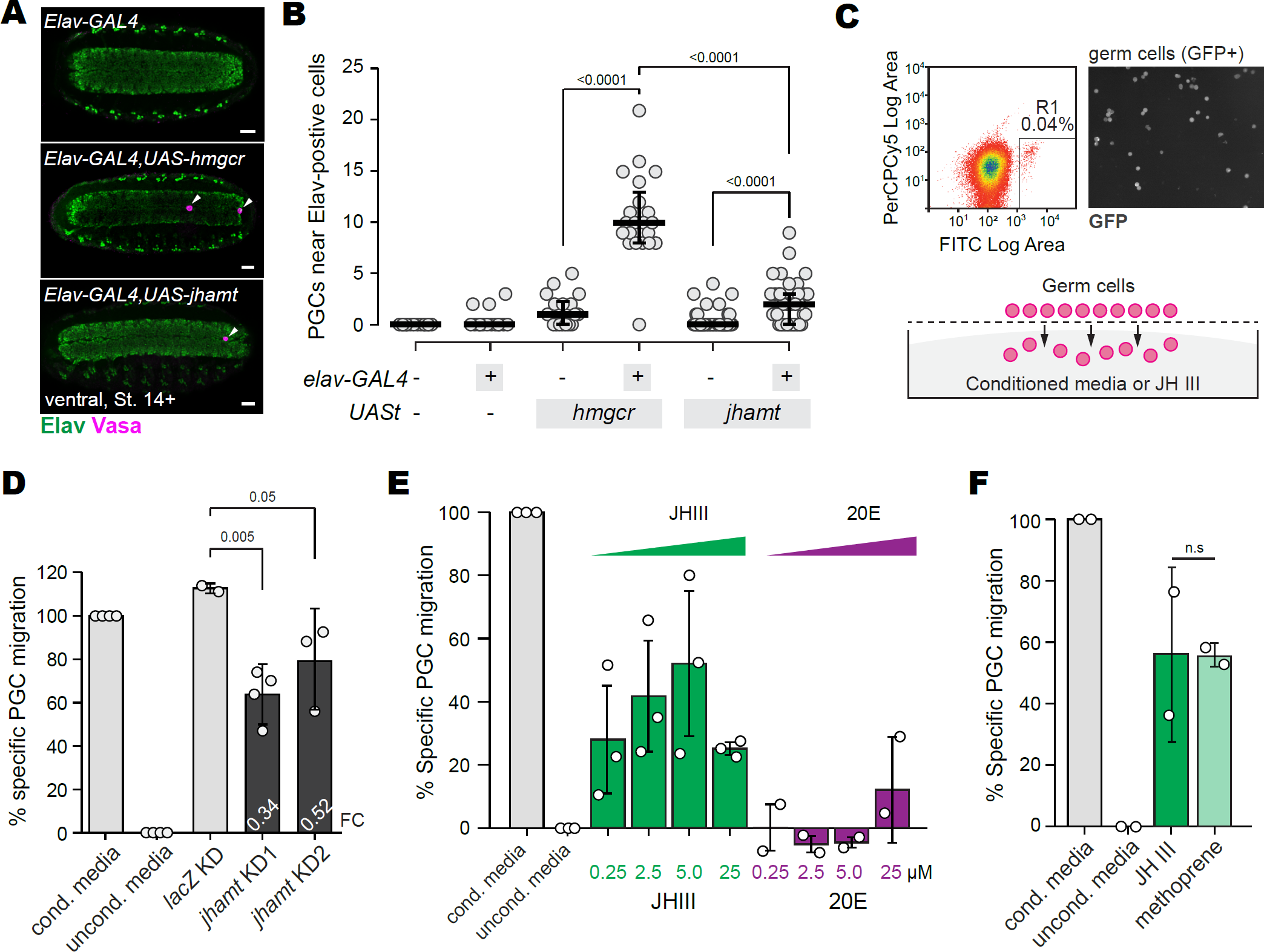
Juvenile hormone is sufficient for PGC migration. **A** Ventral view images of Stage 14 embryos stained for Vasa (magenta) and Elav (green). Genotypes are noted in the top left corner of each image. Arrowheads highlight germ cells in contact with Elav-positive central nervous system. All embryo images are maximum projections of two to three 2µm confocal slices. Scale bars represent 25µm. **B** Quantification of the number of PGCs in Stage 14-16 embryos contacting Elav-positive cells upon ectopic expression of either *Hmgcr* or *Jhamt* using the *Elav-GAL4* driver. Bar and whiskers represent mean +/-standard deviation and statistical significance was tested by unpaired t-test. **C** *Top Left*: Scatter plots from FACS of GFP-positive PGCs from *nos-moesin::GFP* transgenic embryos. Region 1 (R1) denotes collection cutoff. *Top Right*: Image of isolated GFP-positive PGCs (gray). *Bottom*: Schematic of the *in vitro* transwell assay in which FACS-enriched PGCs are placed on a 10µm filter above either serum-free media conditioned with Kc167 cells as a positive control, serum-free unconditioned media as a negative control, or experimental conditions, such as serum-free media with JH III. The number of PGCs that translocated to the bottom well was quantified visually after 1.5 hours. **D** Quantification of Specific PGC migration, which is the percentage of experimental translocated PGCs relative to positive (serum-free Kc167-conditioned media) and negative (serum-free media) controls. Such normalization is required to compare biological replicates done on different days. See Materials and Methods for further details. Numbers shown in white within each *jhamt* knockdown bar represents the Fold Change (FC) of *jhamt* mRNA at the time when conditioned media was collected. Bar height represents mean and error bars represent standard deviation from two to four biological replicates and statistical significance was tested by unpaired t-test. **E** Quantification of specific migration in transwell assays using isolated PGCs placed above serum-free media containing various concentrations of commercially available JH III (green bars) or 20-hydroxyecdysone (20E, magenta bars) with 1% BSA as a non-specific carrier protein. Bars represent means and error bars represent standard deviation from three biological replicates, which were each obtained by averaging two to three technical replicates. **F** Percentage Specific PGC migration wherein isolated PGCs were placed across either serum-free media conditioned (gray), serum-free media with 1% BSA (black) and either JH III (dark green, 2.5µM) or the JH analogue, methoprene, (light green, 2.5µm) with 1% BSA as a non-specific carrier protein. Bars represent means and error bars represent standard deviation from two biological replicates, which were each obtained by averaging two to three technical replicates. ns=not significant.

Hmgcr and Jhamt are both required for JH biosynthesis. There are at least two possible explanations for the difference by which ectopic expression of *jhamt* and Hmgcr was sufficient to attract PGCs *in vivo* (**Supp Fig 3b**). Hmgcr may direct PGCs through production of JH and non-JH signals. Indeed, Hmgcr is required for the synthesis of many molecules and post-translational protein modifications^29^. Alternatively, *jhamt* expression may not be sufficient in every tissue due to a lack of precursors derived from the upstream mevalonate pathway. Indeed, Hmgcr is the rate-limiting enzyme in the mevalonate pathway and it’s expression is spatially restricted during mid-embryogenesis^19^. To assess the role of JH directly, we turned to an *in vitro* transwell migration assay, in which *Drosophila* PGCs were obtained from whole embryos, enriched by fluorescence-activated cell sorting (FACS), and placed on a porous filter above media (**Fig 3c**). Previous work showed that PGCs will translocate in response to serum-free culture media conditioned with Kc167 or S2 embryonic *Drosophila* cells^30^ (**Fig 3d**). To explore the impact of JH on PGC migration in this *in vitro* migration assay, we knocked down *jhamt* mRNA in Kc167 cells using two, independent dsRNAs (**Supp Table 3**), collected conditioned media, and measured PGC translocation. Like Hmgcr depletion^30^, significantly fewer PGCs translocated in response to media that was conditioned by *jhamt*-depleted Kc167 cells compared to control conditioned media (**Fig 3d**). To verify that reduced PGC translocation was not due to fewer Kc167 cells conditioning the media, we quantified Kc167 cell number and found that *jhamt* depletion had no effect (**Supp Fig 3b**). These data indicated that Jhamt, produced in somatic cells, facilitated PGC migration *in vitro*.

We next asked whether JHs are sufficient for PGC migration. To this end, FACS-enriched PGCs were placed on the transwell filter above serum-free, unconditioned culture media supplemented with commercially available JH III with 1% BSA as a non-specific carrier protein. We found that PGCs translocated in response to JH III, with a positive correlation with JH concentration except at the highest JH concentration tested (**Fig 3e**). To test the specificity of this response, we subjected PGCs to the JH III analogue and insecticide, Methoprene, and found that PGCs translocated in response to Methoprene at levels comparable to JH III (**Fig 3f**). Together, these data demonstrated that JHs were sufficient to induce or direct PGC migration.

### Compensatory feedback maintains JH homeostasis in *Drosophila melanogaster*

Our findings suggested that JHs originating from the somatic mesoderm direct PGC migration to the somatic gonad. We next investigated the mechanisms by which PGCs respond to JHs. JH signaling response canonically involves JHs binding to the Met/Gce receptor, which then associates with a co-activator Taiman to influence transcription of target genes (**Fig 1a**)^15^. To determine whether the JH receptors are required for PGC migration, we analyzed embryos mutant for the two, partially redundant, JH receptors^14, 15^. As existing *Met, gce* double mutant strains die during metamorphosis^14^, to test the maternal and zygotic contributions of Met/Gce we generated new *Met*^1^*, gce ^Mi^*^02742^ double mutants. In *gce^Mi^*^02742^*, gce* mRNA is reduced by more than 90% (**Fig 4a** and **b**) and *Met*^1^ carries a point mutation that renders *Drosophila* 33-fold more resistant to lethality inflicted by methoprene^31^. This strain was crossed with *Met*^27^*, gce^2.5k^* heterozygous females. Even under non-crowding conditions, only ∼5% of the expected class of heteroallelic *Met, gce* double mutant animals eclosed. As expected^14, 32^, the fecundity of these escaper *Met, gce* females was significantly reduced (**Fig 4c**), but nevertheless, allowed us to analyze Met and Gce requirements in PGC development.

**Figure 4:**
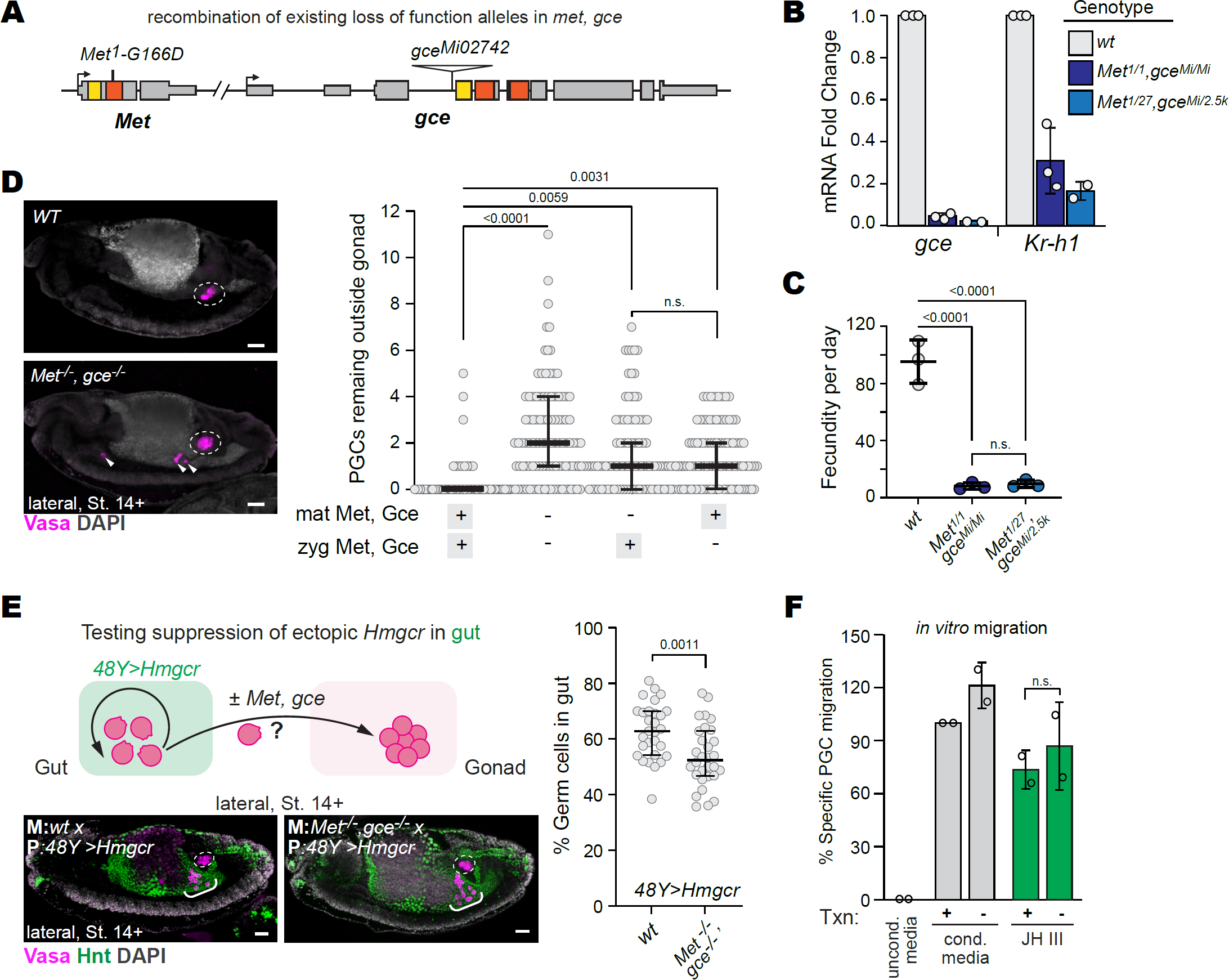
JH transcriptional receptor loss compromises PGC migration to the gonad. **A** Schematic of genetic loci encoding the two redundant JH transcriptional receptors, Met and Gce with location of extant mutations recombined for this study noted. **B** Fold change of *gce* and *Kr-h1* mRNAs in *met, gce* mutant one-day old adult females relative to wildtype control animals. deltaCt was calculated relative to the average Ct of *DCTN5-p25* and *Und*. Shown are Fold Change in homoallelic combination for the newly recombined mutant alleles *Met^1^, gce^Mi^* (dark blue) and heteroallelic combination of *Met^1^, gce^Mi^* with existing or *Met*^27^, *gce^2.5k^* alleles (light blue). Bars and error bars represent mean +/-standard deviation from three biological replicates, each containing pooled mRNA from five females. **C** Fecundity of young females. Each dot represents the average number of eggs laid per biological replicate, which were obtained by dividing the number of eggs laid over a 24-hour period by the number of females (10-12) in each cohort three- and four-days after eclosion. To avoid any confounding male infertility effects, all virgin females were mated to wildtype (*w*^1118^) males. Bars and whiskers represent mean +/-standard deviation and statistical significance was tested by unpaired student t test. **D** *Left*: Lateral view images of Stage 14 embryos stained for Vasa (magenta) and DAPI (gray). Maternal and zygotic genotypes are noted in the top left corner. *Right*: Quantification of extragonadal PGCs in Stage 14-16 embryos. Each dot represents one embryo. To generate maternal and zygotic mutant embryos, mutant *Met^1/27^, gce^Mi/2.5k^* mothers were crossed to *Met^1^, gce^Mi^/Y* males. Arrowheads point to extragonadal germ cells. **E** *Top panel*: schematic of genetic suppression experiment. In otherwise wildtype embryos, ectopic expression of *Hmgcr* in the gut/endoderm (green) with *48Y-GAL4; UASt-Hmgcr* transgenes traps ∼65% of PGCs (pink) in the gut (*left schematic*). To determine if Met and Gce is required downstream of Hmgcr/JH, *Met*^1^*^/^*^27^*, gce^Mi/^*^2^*^.^*^5k^ females were crossed *48Y-GAL4; UASt-Hmgcr* males to generate PGCs without maternal loading of Met/gce. *Bottom panels*: Lateral view of Stage 14 embryos stained for Vasa (magenta), the endodermal marker, Hnt (green) and DAPI (gray). Bold M: designates the maternal genotype that PGCs primarily rely on when exiting the endoderm. Bold P: designates the paternal genotype. White brackets indicate germ cells that remain in gut. *Right*: Percentage of PGCs remaining in the gut in Stage 14+ embryos upon expression of *UAS-Hmgcr* using the *48Y-GAL4* driver. Each dot represents one embryo. **F** Quantification of Specific *in vitro* migration when isolated PGCs were treated with 1µM flavopiridol to inhibit transcription (Txn) during and 30 minutes prior to and during the same transwell migration assay described in Fig 3c-f. Specific migration was calculated relative to vehicle control treated PGCs subjected to serum-free media (uncond media, black) and serum-free media conditioned with Kc167 (Cond media, gray). The effect of transcriptional inhibition (-) was measured in PGCs subjected to cond media or containing 5µM JH III (green). Bars represent means and error bars represent standard deviation from two biological replicates, which were each obtained by averaging two technical replicates. Bars and whiskers within each graph represent median +/-interquartile range (IQR). Statistical significance was tested by Mann-Whitney U test, with ns=not significant. Dashed white circles outline the gonad and scale bars represent 25µm in all images.

To investigate PGC response requirements, we stained maternally and zygotically *Met, gce* mutant embryos for Vasa and DAPI. Embryos lacking either maternal or zygotic Met and gce showed a moderate loss of PGCs relative to wildtype controls at the end of migration (**Fig 4d**). However, we observed a significant increase in the number of extragonadal PGCs in embryos lacking both maternal and zygotic Met and gce (**Fig 4d**). Because PGCs primarily rely on maternally deposited mRNAs during migration^33^, the additive effect of maternal and zygotic loss suggests either an early somatic requirement or a late PGC requirement for Met/gce. To test for a link between JH receptors and Hmgcr, we next asked whether Met/Gce suppressed the ability of PGCs to respond to Hmgcr. For this experiment, we took advantage of our observation that ectopic *Hmgcr* expression in the endoderm trapped 50-80% of PGCs within the gut at Stage 10 (**Fig 4e**). To test whether loss of Met and Gce suppressed the ability of ectopic *Hmgcr* expression to trap PGCs, we crossed wildtype or *Met, gce* double mutant females to males containing *48Y-GAL4* and *UASt-Hmgcr* transgenes and quantified the percentage of PGCs remaining in the gut after gonad colonization (**Fig 4e**). This experiment revealed a small, but statistically significant, suppression of gut trapping in maternally *Met, gce* mutant PGCs (**Fig 4e**). Together, these data are consistent with a requirement for Met and Gce in PGC migration to the gonad downstream of Hmgcr/JH signaling.

Canonical JH signaling leads to transcriptional changes of target genes. To determine whether transcription is required for PGCs to respond to JHs, we returned to the *in vitro* migration assay, where we inhibited transcription specifically in PGCs using the P-TEFb-associated CDK9 inhibitor, flavopiridol. As expected^34^, flavopiridol rapidly and robustly inhibited transcription in PGCs enriched by FACS (**Supp Fig 4**). We then subjected control and transcriptionally deficient PGCs to conditioned media, serum-free media, or serum-free media containing JH III and found that inhibition of transcription did not alter PGC translocation in response to any condition tested (**Fig 4f**). These data suggest that the migratory response of PGCs to JH does not depend on classical transcription-dependent JH receptor signaling.

We were surprised that PGC migration to the gonad was compromised in *Met, gce* mutant animals *in vivo*, yet transcription was not required for JH-mediated PGC migration JHs *in vitro*. However, we noticed that extragonadal PGCs in *Met, gce* mutant embryos were often located centrally and anterior to the gonad, rather than posterior to the gonad as observed in *jhamt* and *Hmgcr* mutant embryos (**Fig 2d** and **Fig 4d**). The central, anterior location of extragonadal PGCs in *Met, gce* mutant embryos suggests that migration to the gonad may be compromised at an earlier stage than in *jhamt* mutant embryos. One possible explanation is that altered JH homeostasis within the embryos, rather than loss of JH signaling response in migrating PGCs, were the cause of extragonadal PGCs in *Met, gce* mutants. To test this possibility, we measured JH titers by quantitative mass spectrometry. Interestingly, we found that both JH III and JHB3 titers were dramatically increased in *Met, gce* mutant animals relative to wildtype (**Fig 5a**). To investigate how JH titers could be increased in *Met, gce* mutant animals, we measured *jhamt* mRNA levels by real time-qPCR. We found increased *jhamt* mRNA levels in *Met, gce* mutant<u>s</u> in several life stages, including during mid-stage embryo embryogenesis (**Fig 5b** and **Supp Fig 5a**). Consistent with compromised JH-dependent transcriptional signaling, expression of the target gene, *Kr-h1*, was not increased in *Met, gce* mutant animals (**Fig 5b**). These data uncovered a compensatory feedback mechanism in *Drosophila melanogaster* by which JH receptors mediate JH titer homeostasis. Importantly, this study suggests that PGC migration to the gonad is compromised by either decreased or increased JH levels.

**Figure 5:**
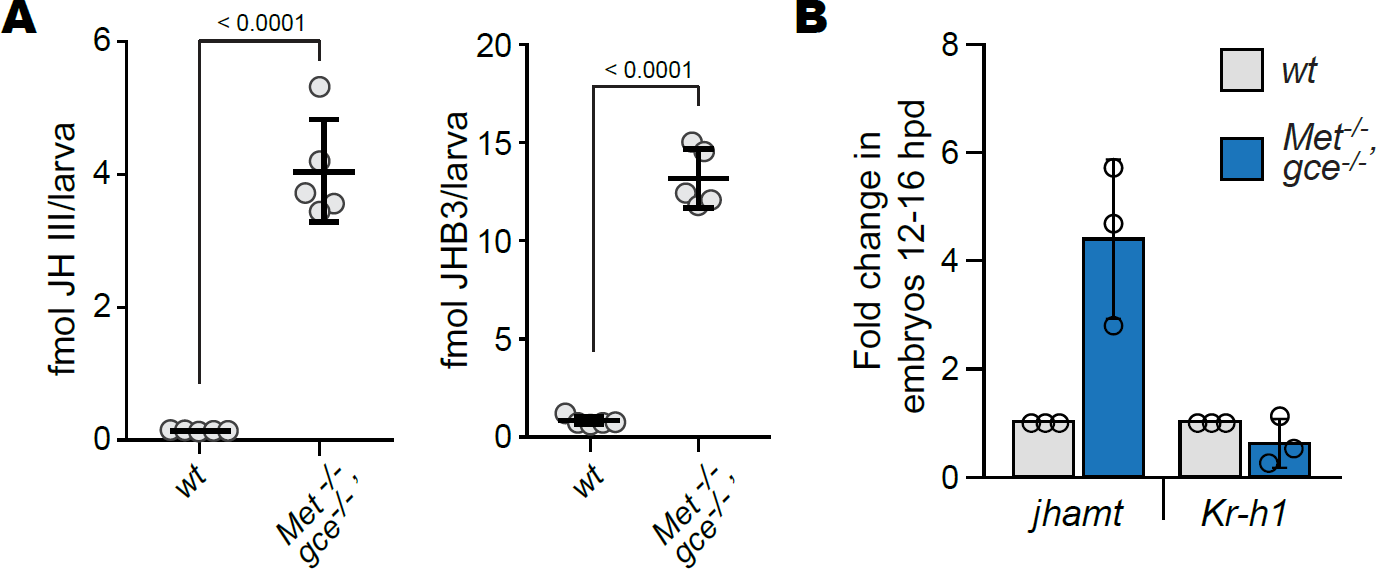
Compensatory feedback loop maintains JH homeostasis in *Drosophila*. **A** Titers of two JHs, JH III and JH bisepoxide (JHB_3_), in hemolymph of third instar *Drosophila* larvae measured by quantitative mass spectrometry using deuterated standards. Each dot represents a biological replicate of hemolymph bled from 25 *Met*/, *gce^Mi/Mi^* or *w*^1118^ third instar larvae. **B** Fold change of *jhamt* and *Kr-h1* mRNA in *Met*^1/27^, *gce^Mi/2.5k^* embryos 12-16 hours post deposition (hpd) relative to the wildtype strain obtained by crossing *w*^1118^ to OregonR. deltaCt was calculated relative to the average Ct of two housekeeping genes, *DCTN5-p25* and *Und*. Bars and error bars represent the average fold change and standard deviation from three biological replicates.

### Isoprenoids in PGC migration

JHs belong to a broader class of secreted isoprenoids that have functions in reproduction throughout the animal kingdom. Such molecules include methyl farnesoate, which has potent impacts on sex differentiation in the crustaceans^6^, and retinoic acids (RAs), which have pleiotropic effects at many stages of vertebrate reproductive development and gametogenesis^35–37^. Although the transcriptional receptors differ, they are promiscuous enough to bind to a broad class of exogenous secreted isoprenoids and other small molecules^38^. To determine whether *in vitro* cross-activation extends to PGC migration, we enriched *Drosophila* PGCs by FACS and subjected them to JH III or two RA species, all-trans retinoic acid (ATRA) or 9-cis RA. Notably, 9-cis RA induced *Drosophila* germ cell translocation to levels comparable of JH III (**Fig 6a**). These data show that RAs can induce *Drosophila* PGC migration, suggesting commonality in response to this class of secreted isoprenoids.

**Figure 6:**
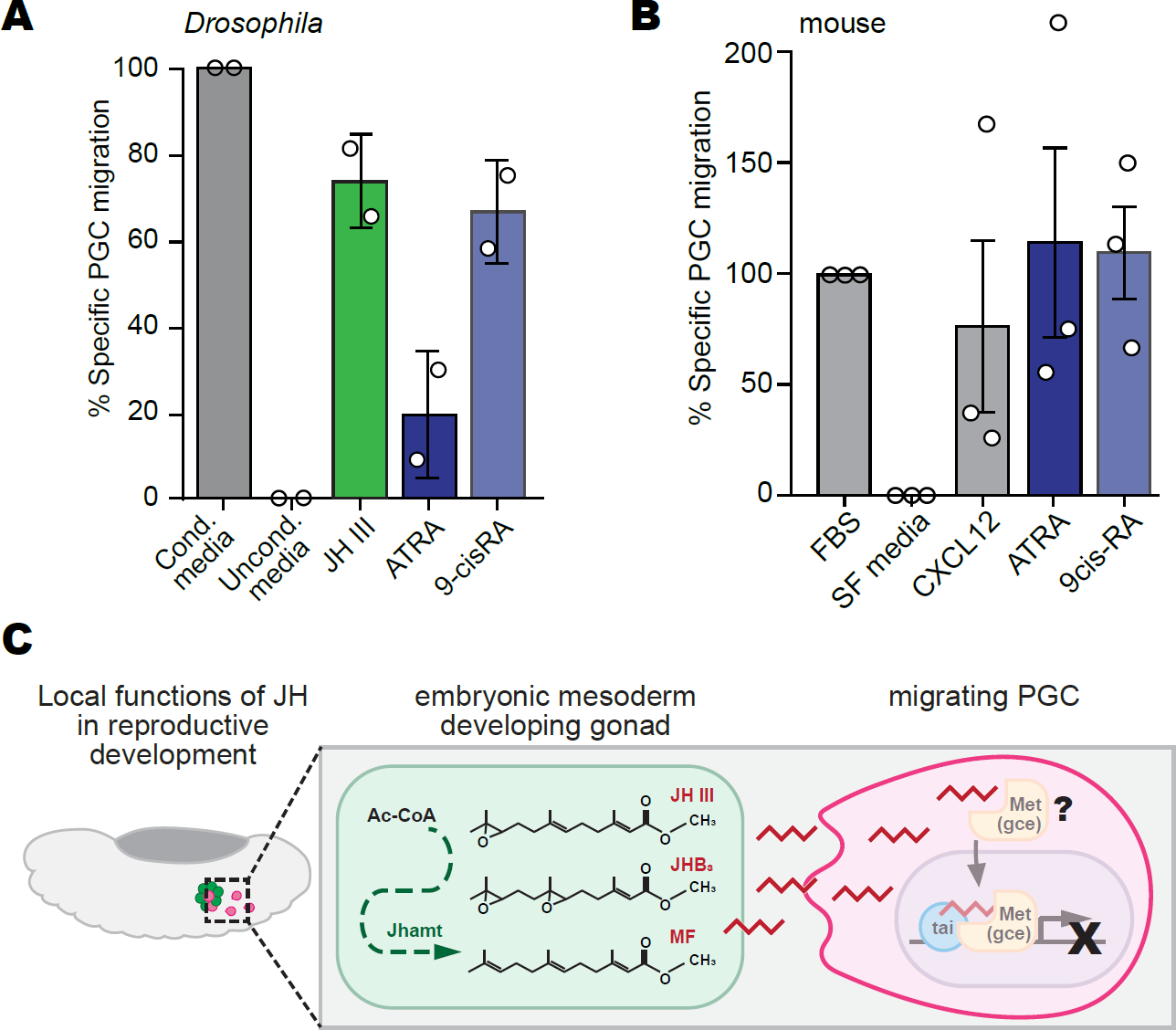
Model of JH function in PGC migration. **A** Quantification of specific *in vitro* migration wherein isolated *Drosophila* PGCs are placed in a transwell migration system across Kc167-conditioned media (gray), serum-free media (black), JH III (green, 5µM), all trans retinoic acid (ATRA, dark blue, 5µM) or 9-cis RA (light blue 5µM). **B** Quantification of specific *in vitro* migration wherein mouse PGCs were FACS-enriched from trunk tissues dissected from E10.5 *Oct4-GFP* embryos and placed across two positive controls: Fetal Bovine Serum (FBS) or CXCL12 (50 ng/mL) (gray), serum-free media as a negative control (black) or ATRA (dark blue, 1µM) or 9cis-RA (light blue, 1µM). For A and B, bars and error bars represent mean +/-standard deviation from at least two biological replicates, which were obtained by averaging two to three technical replicates. **C** Schematic of model for how JHs ensure migrating PGCs (pink) colonize the somatic gonad (green) in the *Drosophila* embryo.

Our studies revealed that the JH biosynthesis enzyme was present in the mesoderm near PGCs as they migrated toward the developing somatic gonad during *Drosophila* embryogenesis. Interestingly, the enzymes critical for RA biosynthesis are also expressed in the developing bipotential somatic genital ridge of mouse embryos as germ cells are colonizing this tissue^21^. To determine whether RAs may have a similar role in vertebrate germ cell development, we adapted the *in vitro* transwell migration assay developed for *Drosophila* PGCs to mouse germ cells. To this end, we obtained mouse germ cells by FACS from embryonic day E10.5 mice carrying a germ cell-specific transgene (*Oct4ΔPE:GFP*) (**Supp Fig 6a**). At this stage in reproductive development, mouse PGCs are completing their migration and beginning to populate the bipotential somatic gonad. We exposed germ cells to fetal bovine serum in PGC media as a positive control or serum-free PGC media as a negative control. Both ATRA and 9-cis RA induced translocation of mouse embryonic germ cells to levels comparable to the known mouse and zebrafish PGC chemoattractant, C-X-C Motif Chemokine Ligand (CXCL12) (also called SDF1a) (**Fig 6b**)^39–41^. Although RAs can have a proliferative effect on mouse PGCs over 24-72 hours^42^, we did not detect changes in total PGC number during our two-hour assay (**Supp Fig 6b**). Therefore, the increased number of translocated PGCs was not due to RA-induced proliferation. Together, these data suggest that the newly uncovered role of secreted isoprenoids in early PGC migration and colonization to the somatic gonad may be broadly shared among invertebrate and vertebrate species.

## Discussion

Germ cells are shepherded through development and differentiation by surrounding somatic cells, which provide hormones and other signaling molecules. Here, we demonstrated that Juvenile Hormones (JHs), key gonadotropins in adult gametogenesis, functioned in embryonic reproductive development. In the embryo, JH signaling was spatially restricted and enriched around PGCs as they migrated to the somatic gonad. We found that JHs were necessary *in vivo* and sufficient *in vitro* for PGC migration. Together, these studies significantly expanded our understanding of the scope, functions, and mechanisms by which JHs support the germline lifecycle.

### JH signaling is active during embryogenesis prior to endocrine gland formation in *Drosophila*

The study of JH biology has largely focused on endocrine production in juvenile and adult animals, particularly for insects that undergo a complete metamorphosis, like *Drosophila*. In juvenile and adult animals, JHs are produced in the CA endocrine gland. CA precursors are first distinguishable at Stage 13 and migrate dorsally to form the ring gland at Stage 15^43, 44^. Through the generation of an *in vivo* JH sensor, we provide evidence that JH signaling is active in *Drosophila* embryos starting at Stage 10 (**Fig 1e** and **f**). Our data are consistent with a burst of *Kr-h1* expression in 8-12 hr old embryos^45, 46^ and with recent reports of non-CA *jhamt* expression in gut cells^47^. Combined, these studies show that sources of these important hormones exist outside of the CA. Three species of biologically active JHs detected in *Drosophila* are proven agonist ligands of Gce^25, 48^, yet whether these forms are temporally and functionally distinct *in vivo* are unknown. The discovery of embryonic JH activity in the genetically tractable *Drosophila* embryo and in an *in vitro* migration assay provide a unique opportunity to explore JH species-specific functions.

### JH signaling is paracrine during mid-embryogenesis

We observed JH activity near mesodermal expression of JH-producing enzyme Jhamt (**Fig 1b, e, f**). This observation suggests that JH signaling is paracrine during embryogenesis. Paracrine signaling mechanisms are consistent with reports that adult gut cells exhibit local JH signaling^49^ and that ectopic *Hmgcr* expression influences PGC migration within a ∼50um zone^28^. How JH signaling response is spatially restricted has yet to be understood, but one possibility is that JH transport or diffusion is limited by the dense extracellular environment of the embryo. Alternatively, JH signaling may be restricted by the spatio-temporal expression of enzymes that synthesize or degrade JHs or be restricted to cells that possess all required factors in the transcriptional signaling cascade. Whether all co-factors are present in all somatic cells during mid-embryogenesis is unknown, however the core JH transcription factors, *Met, gce* and *taiman* mRNAs are maternally loaded into the early embryo^50–53^. Whether JH binding proteins, which facilitate JH circulation in whole juvenile and adult insects, are required for embryonic JH signaling is unknown. The highly dynamic, well-characterized, and pre-circulatory features of mid-embryogenesis can be leveraged to understand the mechanisms of paracrine JH signaling.

### JH signaling ensures migrating PGCs reach the somatic gonad

Embryonic JH signaling is enriched in the trunk mesoderm as PGCs migrate in this tissue toward the somatic gonad (**Fig 1c, e, f**). We found that JH biosynthesis enzymes are necessary for efficient gonad colonization *in vivo* and JHs are sufficient for PGC migration *in vitro* (**Fig 2** and **Fig 3**). While JHs have many functions in reproduction, our findings demonstrate that JH activity impacts germline development far earlier than previously appreciated. Moreover, a role for JHs in cell migration had not been reported to our knowledge. In contrast, ecdysone, the steroid hormone controlled, in part, by JHs^54^, regulates the migration of many somatic ovarian cell types, including swarm cells during juvenile ovarian development^3^ and border cells during adult gametogenesis^4^. Given the antagonistic relationship between the steroid ecdysone and the isoprenoid JHs^54^, it remains possible that JHs also prevent ecdysone-mediated processes while germ cells colonize the gonad. Such regulation by JHs may have been obscured by the relatively high degree of genetic redundancy throughout the JH biosynthetic, degradative, and signaling response cascades in the *Drosophila* genome^15, 25, 26^. The findings described here provide an opportunity to understand the relationship between steroid hormones and JH/retinoid signaling molecules. Indeed, antagonism between these two classes of signaling molecules are observed in multiple tissue contexts throughout the animal kingdom^54, 55^.

Our findings that JHs impact PGC migration opens many questions. Do JHs regulate PGC directionality or motility? What signaling events are required for migrating PGCs to respond to JHs? Classic JH signaling involves receptor-mediated transcription. In contrast, we find that transcription is not required in PGCs for migrational response to JHs *in vitro* (**Fig 4f**). Our finding is consistent with the observation that PGCs respond to ectopic Hmgcr prior to full activation of their zygotic genome (**Fig 4e**) and that migrating PGCs must rapidly respond to environmental changes. Rapid responses to JHs during ovarian follicle development has been observed for decades^56–59^, however the mechanisms remain elusive. Both receptor tyrosine kinases and G protein-coupled receptors (GPCRs) have been implicated in rapid responses to JHs^60, 61^. A requirement for GPCRs is consistent with our observations that migratory response of *Drosophila* PGCs is attenuated at the highest dose of JHs *in vitro* (**Fig 3e**) and compromised when JH titers are elevated *in vivo* (**Fig 4d** and **5a**), a characteristic feature of GPCR desensitization. Identification of a plasma membrane receptor capable of mediating a rapid response to JHs and related molecules during early PGC development will be an exciting area of future exploration.

Signaling responses to JHs usually involve DNA-binding receptors that induce transcription. We found that the *Drosophila melanogaster* JH receptors, Met/Gce, are required for PGCs to efficiently migrate to the gonad (**Fig 4d**) downstream of Hmgcr (**Fig 4e**). However, genetic suppression of Hmgcr-mediated trapping was mild and the location of extragonadal *Met, gce* PGCs differs compared to embryos lacking the JH biosynthesis enzymes. In the absence of Jhamt and Hmgcr, extragonadal PGCs are often found lateral and posterior to the gonads, whereas extragonadal *Met, gce* PGCs are found medially and anterior to the gonads (**Fig 2d** and **Fig 4d**). While anterior extragonadal PGCs could result from failing to stop migrating at the target gonad, the medial localization and frequent endodermal association of *Met, gce* PGCs suggests that migration is compromised earlier than in *jhamt* mutants. There are a few possible explanations for this surprising observation. First, JH signaling may be more compromised earlier and stronger in *Met, gce* mutants than in *jhamt* mutants. Indeed, loss of *jhamt* does not completely abolish JH titers (**Fig 2b**), as has been reported previously^25^. Second, JH signaling may be required maternally as JHs are maternally provided into eggs in some species^62^. However, it should be noted that zygotic loss of Jhamt and Hmgcr is sufficient to compromise PGC migration (**Fig 2d**)^19^. Third, migration may be compromised in *Met, gce* mutant animals because JH titers are increased (**Fig 5a**). Future work will be needed to elucidate precisely how Met and/or Gce are required for PGC development. Yet, our findings suggest that either a specific titer or spatial arrangement of JHs are required for PGCs to efficiently populate the somatic gonad. How PGCs respond to finely tuned JH availability will be an interesting area of future exploration.

### Discovery of a compensatory feedback loop that maintains JH homeostasis

Animals carrying mutations in the JH receptors exhibit dramatically high JH titers (**Fig 5a**). Elevated JH titers coincided with increased *jhamt* mRNA levels at multiple *Drosophila* life stages (**Fig 5a, b** and **Supp Fig 5a**), suggesting that constitutive feedback ensures stage-appropriate JH titers upon genetic perturbations. The mechanism of this compensatory loop is not known, however Met, Ecdysone Receptor (EcR), and their shared co-factor, Taiman, binding has been detected ∼1kb upstream of the *jhamt* transcription start site^63^ (**Supp Fig 5b**). Thus, a simple feedback mechanism may involve either Met/Gce-Tai repression or EcR-Tai activation of *jhamt* transcription. Indeed, reduced *Kr-h1* mRNA observed in *Met, gce* animals (**Fig 5b**) may lead to the upregulation of EcR-dependent signaling as ecdysone biosynthesis genes are repressed by Kr-h1^64, 65^. For these studies, we used the *Met*^1^ allele, which encodes a point mutation in a highly conserved position of the PAS-A domain and the *Met*^27^ allele, which strongly reduces *Met* mRNA but the open reading frame appears wildtype (Marek Jindra, personal communication). Both *Met*^1^ and *Met*^27^ mutant animals have dramatically increased tolerance to methoprene^31^. However, in our hands, adult escapers could only be obtained in heteroallelic *Met*^1^*^/^*^27^*, gce^Mi/^*^2^.^5k^ combination. Even under sparse conditions, homoallelic *Met*^27^*^/^*^27^*, gce*^2^.^5k^*^/^*^2^.^5k^ were not observed, suggesting either the presence of a second site lethal mutation in the *Met*^27^*, gce*^2^.^5k^ genetic background or that *Met*^1^*^/^*^27^*, gce^Mi/^*^2^.^5k^ animals have enough JH signaling to support development through increased JH titers. The later explanation is more likely because UAS-driven *Met* or *gce* can rescue *Met*^27^*^/^*^27^*, gce*^2^.^5k^*^/^*^2^.^5k^ lethality^15^. Moving forward, it will be important to expand the experimental tools available to dissect this robust signaling pathway.

### New animal model to understand isoprenoid signaling in embryonic reproductive development

We found that JH ensures that migrating PGCs reach the somatic gonad of the *Drosophila* embryo. In contrast to systemic endocrine signaling in other life stages, our findings demonstrate that during mid-embryogenesis, JH signaling is active near mesodermal expression of the key JH biosynthesis enzyme, Jhamt. This local, paracrine signaling is similar to how retinoic acids (RAs) act within vertebrate species. Indeed, the bioavailability of RAs and JHs during analogous stages of germ cell development is strikingly similar between mice and *Drosophila*, respectively^20–22, 66^. RAs impact the migration of many cell types. Although most investigations into the role of RAs in cell migration have focused on transcriptional upregulation of chemokines or their receptors^67, 68^, evidence for transcription-independent functions of RAs is an emerging field^69^. RAs facilitate axon guidance in *Xenopus laevis*^70^, and in snails through non-transcriptional mechanisms involving calcium flux, Rho GTPases and G proteins^71–74^. The many similarities in structure, function, and periodic presence throughout the germline lifecycle support the hypothesis that the isoprenoid signaling molecules, JHs and RAs, are functionally analogous in reproductive development across the animal kingdom. We assert that *Drosophila* can be a powerful genetic model to understand the primary functions and molecular mechanisms by which environmentally pervasive isoprenoids impact reproductive development. Such investigations will increase our understanding of how germ cell development is supported by surrounding somatic cells, both endogenously and through *in vitro* gametogenesis technologies.

## Acknowledgements

We thank Marek Jindra and Ryusuke Niwa for reagents and valuable discussions, and Peter Nicholls and David Page for instructive discussions on mammalian reproductive development and assay development. We are grateful to Alexey A Soshnev for graphical designs, Cesar Ramirez and Francisco Fernandez-Lima for assistance with mass spectrometry. We are especially grateful to Rebecca Spokony, Jessica Treisman, Pamela K Geyer, and Toby Lieber for valuable advice and insightful discussions, as well as all members of the Lehmann laboratory for valuable input. Stocks obtained from the Bloomington Drosophila Stock Center (NIH P400D018537) were used in this study. In addition, several monoclonal antibodies used in this study were obtained from the Developmental Studies Hybridoma Bank, created by the NICHD of the NIH and maintained at The University of Iowa, Department of Biology, Iowa City, IA. We, like most users of the *Drosophila* model organism, could not have done this work without the efforts of the FlyBase Consortium (supported by NHGRI #U41HG000739 and #S454486, NSF #2039324, MRC #MR/W024233/1)^75^. We also thank the NYULMC Core Cytometry Facility, which is partially funded by NIH/NCI P30CA016087. This work was supported by a Damon Runyon Cancer Research Foundation postdoctoral fellowship from the William Raveis Charitable Fund (DRG2235-15, L.J.B), an NIH Pathway to Independence award (K99 and R00 HD097306, L.J.B.), an NIH-NIAID R21 award (R21AI167849, F.G.N), and project 22-21244S from the Czech Science Foundation, Czech Republic to MN and an NIH MERIT award (R37HD41900, R.L). RL was a HHMI investigator.

## Author Contributions

Conceptualization: LJB, RL; Experimentation: LJB, JS, EPD MN, FGN, MS; Analysis: LJB, MN, FGN; Writing: LJB, RL; Funding Acquisition: LJB, RL; Supervision: LJB, MS, RL

## STAR Methods

### Animal Maintenance

*Drosophila melanogaster* were raised on medium containing 1.5% yeast, 3.6% molasses, 3.6% cornmeal, 0.112% Tegosept, 1.12% alcohol and 0.38% propionic acid. For embryo collections, apple juice plates contained 25% apple juice, 2.5% sucrose, 2.25% bactoagar, 0.15% Tegosept. Animals were kept at 25°C for all experiments. Fly strains used in this study are listed in Supplementary Table 1.

Mice expressing a germ cell-specific *Oct4τι.PE:GFP* transgene^76^ (B6; CBA-Tg(Pou5f1-EGFP)2Mnn/J) were obtained from Jackson Laboratory (#004654, Bar Harbor, ME, USA). All protocols were approved by the NYU Institutional Animal Care and Use Committee.

### Generation and validation of the P [*JHRE-GFP, w+]* transgenic biosensor

Eight copies of the wildtype JH response element (JHRE) from the *early trypsin* gene from *Aedes aegypti*^23^, or eight tandem copies of the mutant JHRE (Supp Fig 1), were excised from *pGL4.17* vectors provided by Jindra Marek. JHREs were inserted upstream of the minimal *hsp70b* promoter and *eGFP* coding sequence of pS3aG transformation vector (Addgene #31171) containing an *attb* for site-specific integration. JHRE::GFP sensor constructs were integrated into *y+, attP2* at cytological position 68A4 sites by Bestgene.

To validate the JH sensor, transgenes were placed in *met*^1^*, gce^MI^* mutant background. Embryonic GFP signal was measured by immunohistochemistry. To measure JH responsiveness, first instar larvae were placed in food containing 1.6µg of Methoprene per gram of food ^77^, or ethanol as a control for 24 hours. *GFP* and *Kr-h1* mRNA was measured in whole second instar larvae by qPCR using primers in Table 3 (Fig 1 and Supp Fig 1).

### Generation of *jhamt* alleles

The CRISPR guide CCTGGATGTGGTTCAGGATCT was cloned into pU6-2 gRNA vector (DGRC #1363) and injected into embryos into a stock derived from BL51324 carrying *w*^1118^;; *PBacy[mDint2]=vas-cas9VK00027* by Bestgene. Of 29 fertile founders, 87 F1 crosses were set up and screened for *jhamt* mutations by PCR amplification and Sanger sequencing using primers listed in Table 3. Of 48 F1 lines screened, 13 lines were obtained wherein small indels created a frame shift just before the methyltransferase domain, which starts at amino acid 40, leading to a stop codon just downstream, two lines at amino acid 46 (12F1 and 12F2) and the remaining lines at amino acid 54 (1F1, 1F2, 4F1, 4F3, 6F3, 7F2, 7F3, 8F1, 8F2, 12F3, 14M2, 14M3). The effect of new *jhamt* mutations on JH III and JHB_3_ titers were quantified in third instar larval hemolymph.

### Embryo Immunofluorescence

Prior to embryo collection, young parental strains (less than 5 days old) were mated in vials for one day before transferring to embryo collection cages containing apple juice plates and wet yeast. Embryos were dechorionated in 50% bleach for three minutes, rinsed with water and fixed in a scintillation vial containing 2ml of 4% formaldehyde in PBS and 8ml of heptane.

Embryos were fixed for 20 minutes on a shaker. To remove the vitelline membrane, formaldehyde was removed from the scintillation vial and 10ml methanol was added, followed by one minute of vigorous shaking by hand. Embryos were washed with fresh methanol three times before storage at -20°C or future processing.

For immunohistochemistry, embryos were sequentially rehydrated with increasing percentage of PBSTx (PBS with 0.1% TritonX100) and then washed six times with PBSTx before a one-hour block using 1% BSA in PBSTx at room temperature. Primary antibodies were incubated overnight at 4C and then washed six times in PBSTx. Secondary antibodies in 1% BSA in PBSTx were incubated for two hours at room temperature, washed six times in PBSTx. Embryos were then equilibrated in Vectashield (Vector Laboratories, H-1200) overnight at 4°C before mounting on coverslips.

Fluorescent *in situ* hybridization was done as previously shown^78^. Probes were generated using *Drosophila* clones from DGRC (*jhamt* #11840, clone AT13581) and (*Nano*s #1124533, clone RE53469).

### mRNA quantification by qPCR

RNA was obtained by TRIzol (Invitrogen #15596018) extraction. DNA was removed using RQ1 RNase-Free Dnase (Promega #M6101). cDNA was generated using SuperScript III Reverse Transcriptase (Fisher Scientific #18080-044) using both oligo dT and random hexamers. Primers used are listed in Table 3. qPCR was carried out using LightCycler 480 SYBR Green I Master 2X (Roche #04887352001) and a Roche LightCycler 480. Results were calculated as previously described^79^ from crossing points and normalized to one of three reference genes found to have invariant expression across tissues and developmental stages: *DCTN2-p50, und, DCTN5-p25*^80^.

### Evaluation of JH titers by quantitative mass spectrometry

The following genetic backgrounds were used to measure JH III and JHB_3_ titers: *jhamt^8F^*^2^*^/12F^*^1^ paired with the *w*^1118^*; CyO, ft-lacZ* genotype, which was the genetic background the crispr alleles were extensively crossed into and *met*^1^, *gce^Mi^* paired with *w*^1118^. Twenty-five third instar larvae per biological replicate were collected from sparsely populated vials, washed in PBS before placing in 50µl of pre-chilled PBS for hemolymph collection. To collect hemolymph, the cuticle was pinched and pulled apart down the length of the animal. Care was taken not to rupture fat body or intestines. Hemolymph was transferred to pre-chilled silanized glass vials (Fisher Scientific #C4011S5 and #C401154B) using silanized glass pipettes (Brain Bits #FPP).

The glass collection well was rinsed with 50µl additional, pre-chilled PBS to obtain any remaining hemolymph and additional PBS was added to silanized glass vials for a total volume of 150µl. Deuterated JH III (62.5pg) dissolved in acetonitrile (Thermo Fisher #51101) was added to each replicate, then 600µl hexane (Sigma Aldrich #1037011000). Samples were vortexed for 1 minute and centrifuged for 5 min at 2000g at 4°C. Organic phase was transferred to new silanized vial for analysis by mass spectrometry.

For mass spectrometry analysis, the JH III and JH III bisepoxide (JHB_3_) amounts present in hemolymph were quantified by liquid chromatography coupled to tandem mass spectrometry (HPLC-MS/MS)^24^. The identification and quantification of JH III and JHB_3_ in hemolymph samples was based on multiple reaction monitoring (MRM), using the two most abundant fragmentation transitions: for JH III: 267->235 (primary) and 267->147 (secondary), and for JHB_3_: 283->233 (primary) and 283->251 (secondary)^24^.

### Fecundity measurements

Fecundity was measured as previously described^81^ and reported as the average number of eggs laid in three- and four-day old females at 25°C. In each biological replicate, 10-12 females were crossed to 5-6 *w*^1118^ males to mitigate male fertility defects upon *jhamt* or *met, gce* loss.

### Knockdown of *jhamt* in Kc167 Cells

To effectively knockdown *jhamt* mRNA, genomic DNA obtained from Kc167 cells using Qiagen’s Blood and Tissue Kit (#69506) was used as a template. To target *lacZ*, pBluescript SK plasmid used as a template. dsDNA was generated by Choice Taq Mastermix (Denville Scientific #CB4070-8). Gradient PCR was used to obtain optimal dsDNA templates. dsRNAs were generated from dsDNA templates using T7 MEGAScript (Invitrogen #am1334) according to standard protocol. dsRNAs were DNAse treated (Qiagen #79254) and purified using Qiagen RNAeasy Mini Kit (Qiagen #74104) according to instructions. For transfection, Kc167 cells were grown to confluency, resuspended in fresh Schneider’s media (Fisher #21720-001) supplemented with Penn/Strep (Fisher, #15140122) and 10% heat inactivated FBS (Thermo Fisher #16140071) to 1×10^6 cells/ml and allowed to settle overnight. 2µg of dsRNAs were used in conjunction with Effectene Transfection Reagent (Qiagen #301425) into each 6-well dish. After 48 hours at 25°C, media was removed, transfected Kc167 cells were washed in PBS and resuspended in serum-free Schneider’s Media (with Penn/Strep) for three additional days at 25°C prior to collection of Kc167-conditioned media. To determine whether *jhamt* knockdown compromised Kc167 viability, the number of Kc167 cells was counted on the day conditioned media was collected. Knockdown efficiency was determined in Kc167 cells by qPCR when conditioned media was collected for the transwell assay. Primers used to measure knockdown efficiency were chosen outside of region targeted by dsRNAs.

### Isolation of PGCs

*Drosophila* embryos carrying the *P[nos-Moe::eGFP.nos 3’UTR]* transgene from an overnight collection were dechorionated in 50% bleach for two minutes, rinsed and placed in filter-sterilized Chan and Gehring’s Balanced Saline, PGC Sort Buffer (3.2g/L NaCl, 3.0g/L KCl, 0.69g/L CaCl2.2H2O, 3.7g/L MgSO4.7H2O, 1.79g/L Tricine buffer (pH 7), 3.6g/L glucose, 17.1g/L sucrose, 1g/L BSA, fraction V). Dechorionated embryos were manually homogenized in PGC Sort Buffer with Dounce homogenizer (Pyrex #7727-15) and passed through a 100µm filter and 20µm filter and placed in pre-chilled polypropylene falcon tubes (#352063). Live, GFP-positive *Drosophila* PGCs were sorted using a MoFlo XDP cell sorter fitted with a 70µm nozzle into Maxymum recovery microtubes (Axygen #311-05-051) containing 200µl Schneider’s Media.

To enrich for mouse PGCs by FACS, B6; CBA-Tg(Pou5f1-EGFP)2Mnn/J (Jax #004654) mice were bred to produce embryos homozygous for the *Octý*4PE::GFP transgene. Embryonic day 0.5 (E0.5) was assumed to be noon of the day the vaginal plug was observed. Pregnant females were euthanized by carbon dioxide followed by cervical dislocation. Trunks were dissected from E10.5 embryos in chilled PBS. For each two embryos, tissue was dissociated in 300µl of 0.5% trypsin (0.25%, Life Technologies #25200056) for 15 min at 37°C, with occasional manual disruption by pipette. Trypsin was inactivated with 200µl of mouse PGC culture media [DMEM/F-12 (Thermo Fisher #11-320-033), 250µM sodium pyruvate (Thermo Fisher #11360070), 2mM L-glutamate (Gibco #25030081), 1X MEM Non-essential amino acids (Thermo Fisher #11-140-050) containing 25% Bovine Albumin Fraction V (Thermo Fisher #15260037). Dissociated tissue was centrifuged at 200g for 7 min and cells were resuspended in 2mL of PGC media. Cell suspension was filtered through a 35µm cell strainer into a 5 ml falcon round bottom tube (VWR #21008-948) for sorting using a MoFlo XDP cell sorter fitted with a 100µm nozzle. Live, GFP+ germ cells were sorted into Maxymum recovery microtubes (Axygen #311-05-051) containing mouse PGC media.

### *in vitro* Migration Assays

For *in vitro* transwell assays, all plastic ware, tips, tubes and transwell plates used throughout the assay were pre-coated with PBS containing 1% BSA to prevent hydrophobic molecules from sticking to the surfaces. Assays with light sensitive isoprenoids were done in dark conditions as much as possible. For Kc167-conditioned media, serum-free media was added to washed, confluent Kc167 cells and collected three days later. Conditioned media was collected and centrifuged 4,000rpm for 20 min to remove all cellular debris and placed on ice. All isoprenoids (Supplementary Table 4) were prepared by dilution series in either serum-free Schneider’s media for *Drosophila* PGC assays or mouse PGC media. Transwell assays were done using the 96-well ChemoTx Disposable Chemotaxis System (NeuroProbe #116-10) which had a standard PCTE filter membrane with 10µm pore size, 300µl well capacity and 5.7mm cell site diameter. Sorted germ cells recovered on ice for 30 min while the transwell system was set up. Sorted germ cells were then diluted 10-15-fold in serum-free Schneider’s media for *Drosophila* PGC assays or mouse PGC media so that 60µl diluted PGCs were placed above each well. Transwell plates with *Drosophila* PGCs were incubated 25°C for 1.5 hours and those with mouse PGCs were incubated at 37°C for 2 hours. After the incubation period, PGCs were removed from above membrane, the plate was centrifuged at 460rpm for 10min to dislodge cells from the well side of the membrane. Translocated germ cells were removed from the 96-well plate by pipetting up and down and transferring to Nunc Lab-Tek II Chambered Coverglass (Thermo Fisher #155409) for manual quantification by fluorescent microscopy.

The number of PGCs in each top well was calculated by setting aside 10% input (6µl) in duplicate into Lab-Tek wells, each with 300µl serum-free media at the beginning of the transwell assay and quantifying the number of germ cells when the experimental translocated germ cells were quantified. To compare biological replicates done on different days, Specific Migration was calculated by first subtracting the number of germ cells that translocated in the negative control wells (serum-free media) and then calculating the percentage of normalized experimental condition to the respective positive control. Each transwell assay is the result of two to four biological replicates, each consisting of two technical replicates.

### Verification of Hormone Activity *in vitro*

To verify the activity of JH III and 20-hydroxecdysone (Supplemental Table 4), hormones were added to 50% confluent Schneider 2 (S2) cells at the concentrations indicated in Supp Fig 3c. After a six-hour incubation at 25°C, S2 cells were harvested, and mRNA was extracted and analyzed using protocols described in the ‘*mRNA quantification by qPCR*’ Methods subsection.

### Transcriptional Inhibition in *Drosophila* PGCs

Efficient inhibition of transcription in *Drosophila* PGCs isolated by FACS was determined by testing different concentrations of Actinomycin D and Flavopiridol (See Supplemental Table 4). To visualize transcription by immunohistochemistry, Lab-Tek II 8-chambered coverglass were incubated with poly-L-Lysine solution (Sigma-Aldrich #P4707) for 30 min at room temperature and then dried for 1.5 hours. Sorted *Drosophila* PGCs were added to chambers and incubated with Actinomycin D or Flavopiridol for either 15 min or 3 hours, then fixed with 4% paraformaldehyde for 15 minutes, washed in 0.05% PBS with TritonX100, blocked in 1% BSA for 30 min and stained for Vasa, RNA Pol II Ser2CTD and RNA Pol II Ser5 CTD (See Supplemental Table 2 for antibody information). As expected^34^, Flavopiridol rapidly inhibited transcription in *Drosophila* PGC while not compromising cell viability. For transwell assays, sorted *Drosophila* PGCs were first incubated with 1µM Flavopiridol for 30 min prior to placing on transwell membrane and solutions in the bottom well chambers also contained 1µM Flavopiridol to ensure PGC transcription was inhibited throughout the assay.

### Microscopy and Image Processing

Most embryos and other samples were imaged with a Zeiss LSM800, an upright laser scanning confocal microscope using Zen Blue 2.3 software and a Plan-Apochromat 20× 0.8 NA air objective and 40× 1.3NA oil objectives with pinhole size for all imagine was set to 1AU. Images were processed using ImageJ/FIJI.

### Statistics

Statistical analysis was done as described in the Figure Legends. Briefly, for quantification of phenotypic data, such as extragonadal PGC quantification, data was not always normally distributed and thus median +/-IQR are reported, and statistical significance was assessed by Mann-Whitney U test. For mRNA Fold Change and *in vitro* Specific Migration results, mean +/-standard deviation is reported and statistical significance was tested by unpaired t-test. Sample size per biological replicate was influenced by power analysis based on previously observed phenotypic variation.

---

**Supplemental Table 1:**
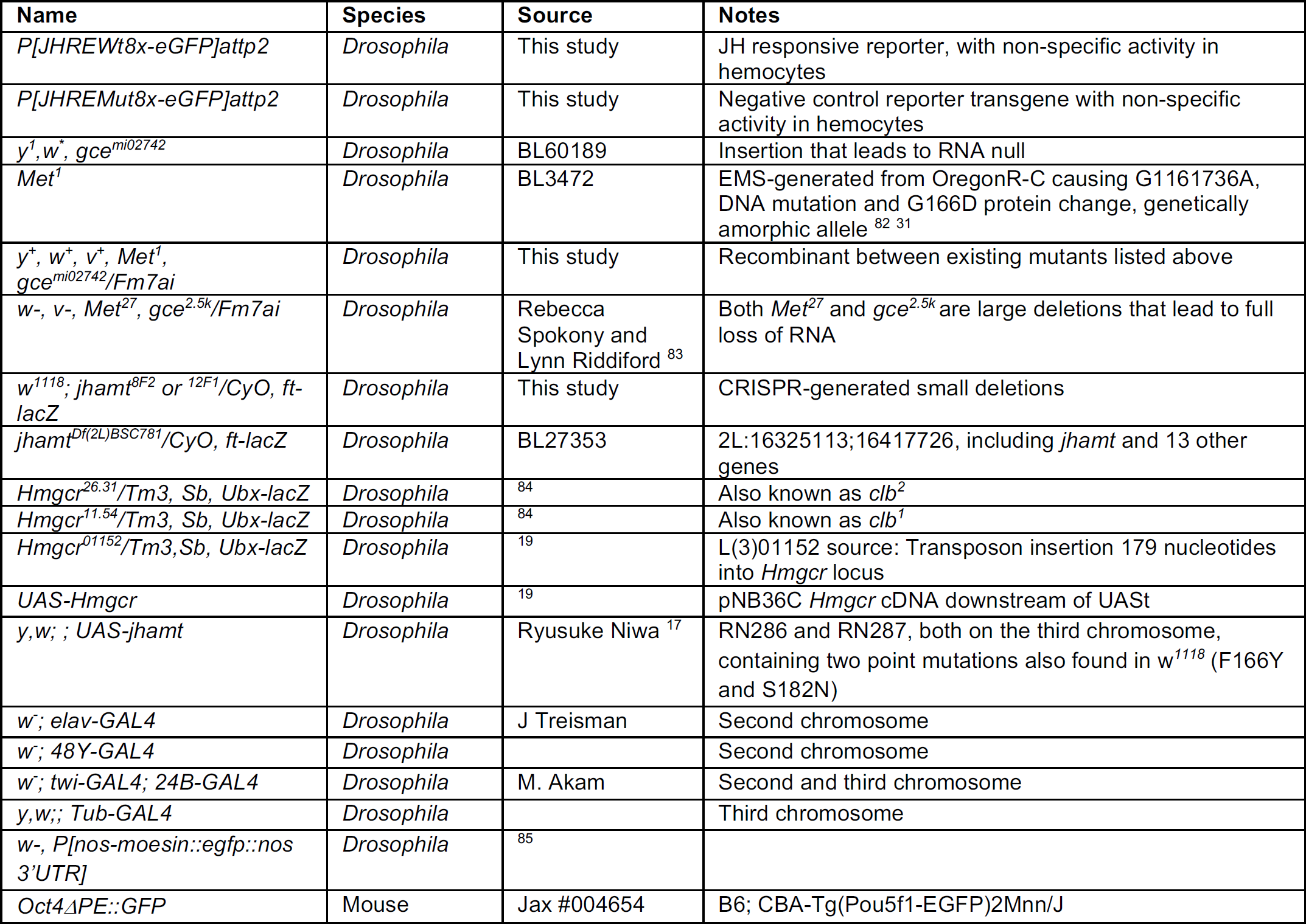
Animal lines used in this study

**Supplemental Table 2:**
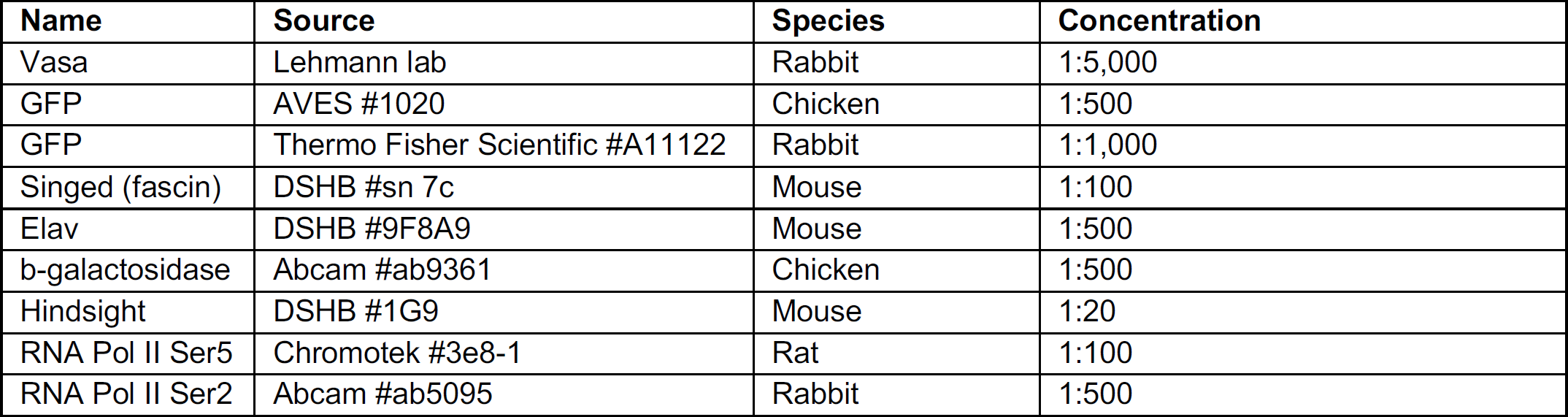
Antibodies used in this study

| Name | Source | Species | Concentration |
| --- | --- | --- | --- |
| Vasa | Lehmann lab | Rabbit | 1:5,000 |
| GFP | AVES #1020 | Chicken | 1:500 |
| GFP | Thermo Fisher Scientific #A11122 | Rabbit | 1:1,000 |
| Singed (fascin) | DSHB #sn 7c | Mouse | 1:100 |
| Elav | DSHB #9F8A9 | Mouse | 1:500 |
| b-galactosidase | Abcam #ab9361 | Chicken | 1:500 |
| Hindsight | DSHB #1G9 | Mouse | 1:20 |
| RNA Pol II Ser5 | Chromotek #3e8-1 | Rat | 1:100 |
| RNA Pol II Ser2 | Abcam #ab5095 | Rabbit | 1:500 |

**Supplemental Table 3:**
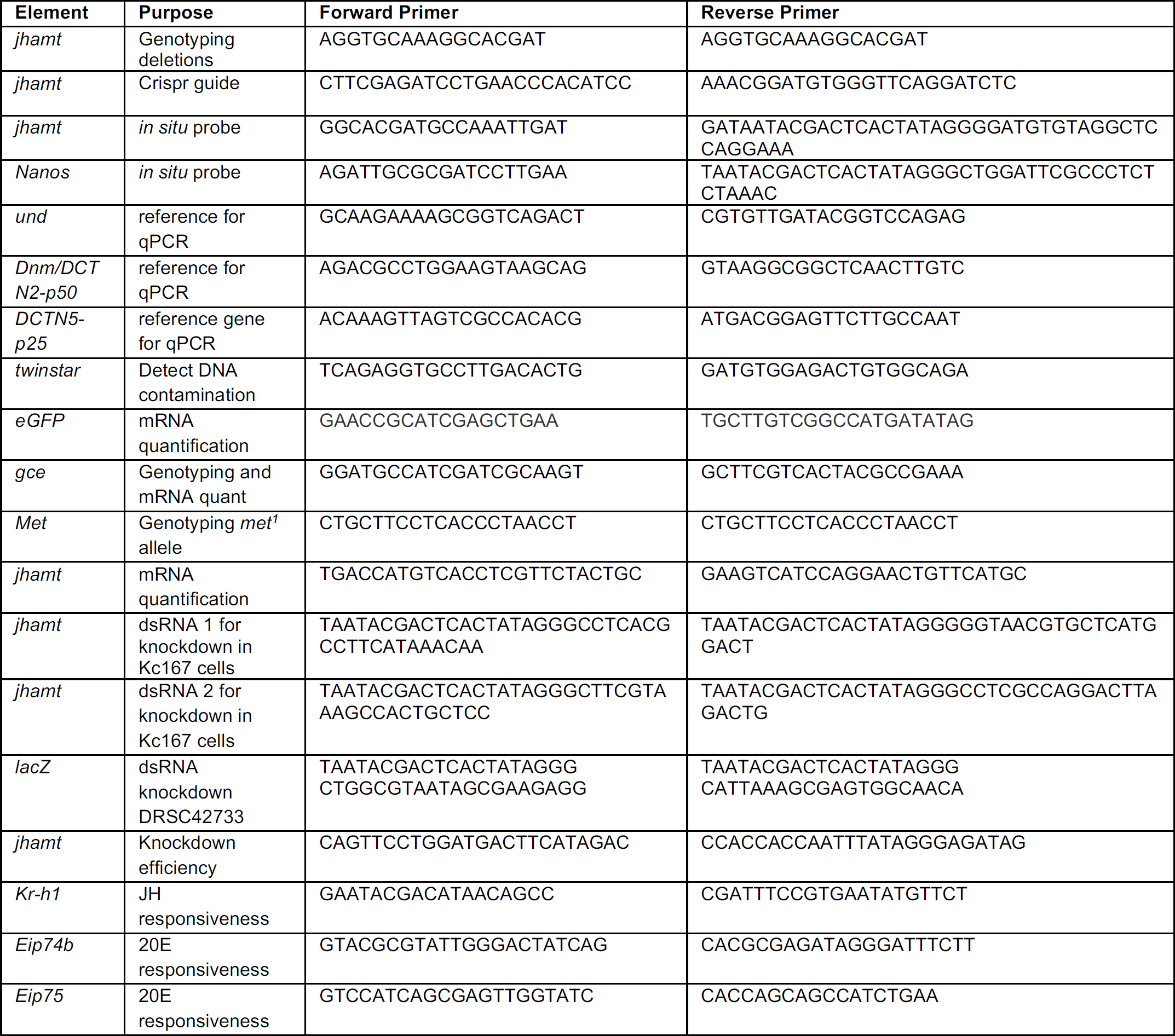
Primers used in this study

**Table 4:** Reagents used in this study

| Name | Source | Product Number | Stock solvent | Final Concentration |
| --- | --- | --- | --- | --- |
| JH III | Sigma-Aldrich | J2000-10MG | Ethanol | 0.25-25µM |
| Methoprene | Sigma-Aldrich | 33375-100MG | Ethanol | 2.5µM |
| 20-hydroxecdysone | Sigma-Aldrich | H5142-5MG | Ethanol | 0.25-25µM |
| ATRA | Sigma-Aldrich | R2625-50MG | DMSO | 2.5µM |
| 9cis-RA | Sigma-Aldrich | R4643 | DMSO | 2.5µM |
| Flavopiridol | Selleck | S1230 | PBS with 0.1% BSA | 1µM |
| Actinomycin D | Sigma-Aldrich | A1410-Smg | DMSO | 1-5µg/mL |
| CXCL12/Sdf-1a | R&D Systems | 460-SD-010 | PBS with 0.1% BSA | 50ng/mL |

**Supplemental Data for Figure 1.**
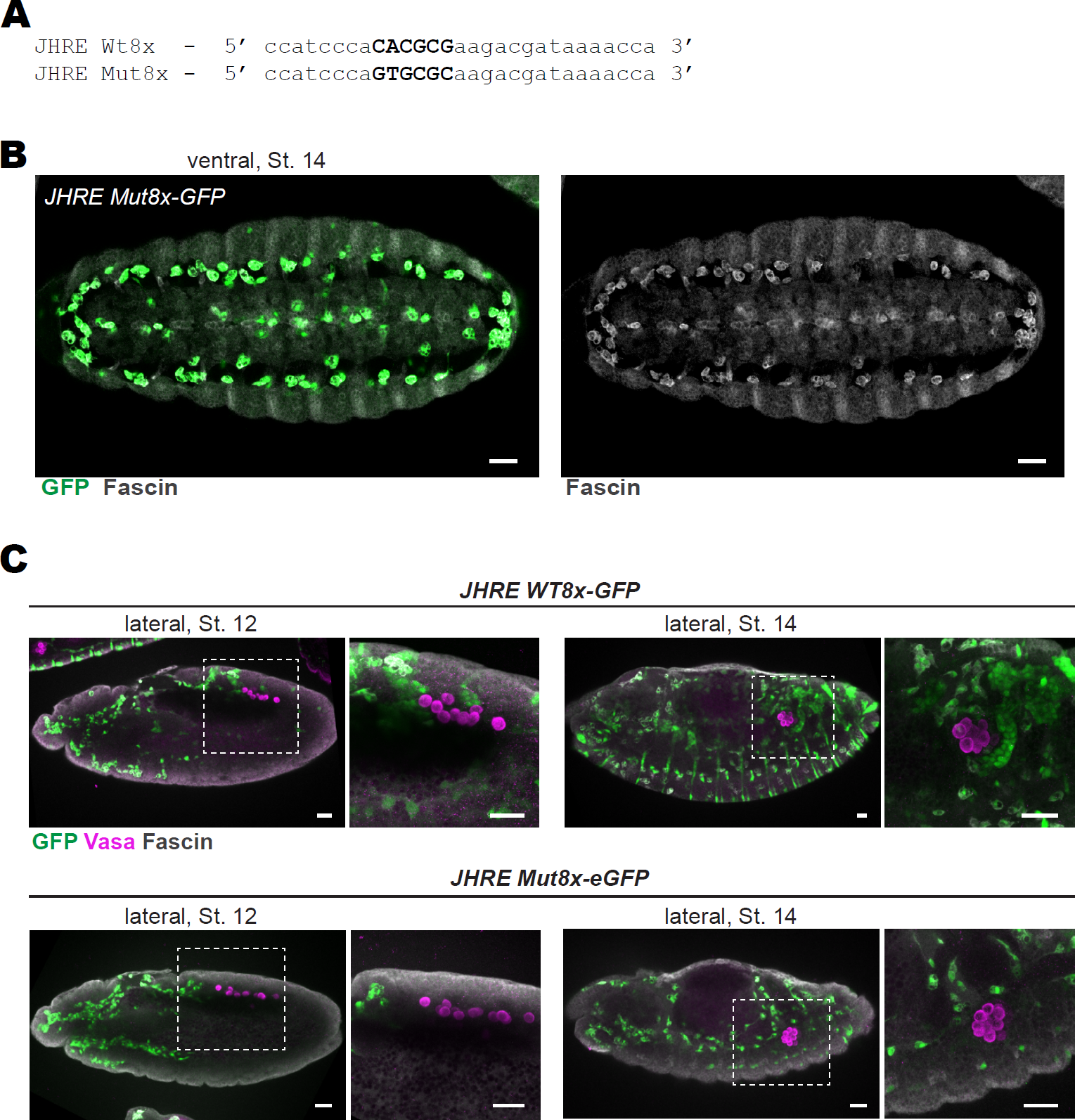
**A** Sequence of the JH response element (JHRE) from the *early trypsin* gene of *A. aegypti* that contains the JH transcription factor, Met/Gce binding motif highlighted in bold^23^. The six base pair wildtype JHRE sequence is on top and the JHRE in which the Met/Gce-binding motif was mutated is on the bottom. Eight copies of each sequence were provided by Marek Jindra^15^ and placed upstream of *eGFP*. **B** Ventral view of Stage 14 embryos carrying a mutated JH biosensor (*JHRE Mut8x-GFP*) in which the JHRE has been mutated. Embryos were stained for GFP (green) and the hemocyte marker, Fascin (gray). Right panel is the same image but with the Fascin signal alone in grayscale. **C** Lateral view of embryos stained for Vasa (magenta), GFP (green) and DAPI (gray). Genotype is noted above each early Stage 12 and Stage 14 pair. To the right of each whole-embryo image is the area within the white, dashed box magnified. Scale bars represent 25µm in all images.

**Supplemental Data for Figure 2.**
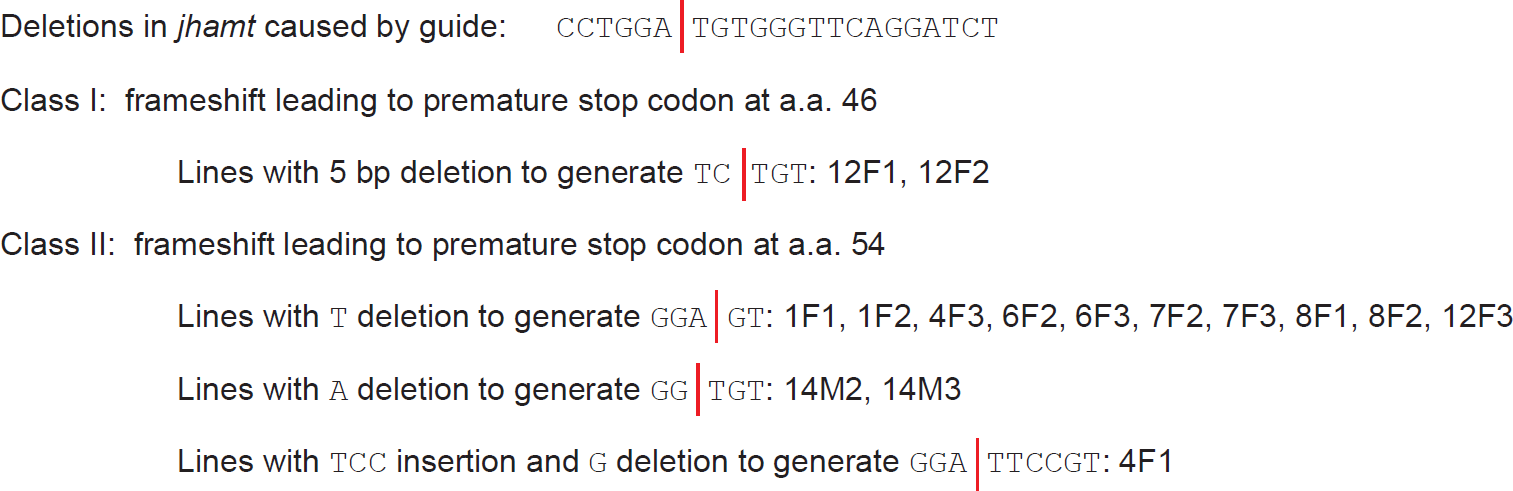
Nature of the small deletions in *jhamt* created by CRISPR-mediated cutting. Seven founders generated fifteen lines which possessed small insertions or deletions at the cut site that created premature stop codons at the beginning of methyl transferase domain of *jhamt* (amino acids 40-137). Class 1 lines (12F1, 12F2) cause a premature stop codon at amino acid 46, while Class II mutations (1F1, 1F2, 4F1, 4F3, 6F2, 6F3, 7F2, 7F3, 8F1, 8F2, 12F3, 14M2, 14M3) cause a premature stop codon at amino acid 54.

**Supplemental Data for Figure 3.**
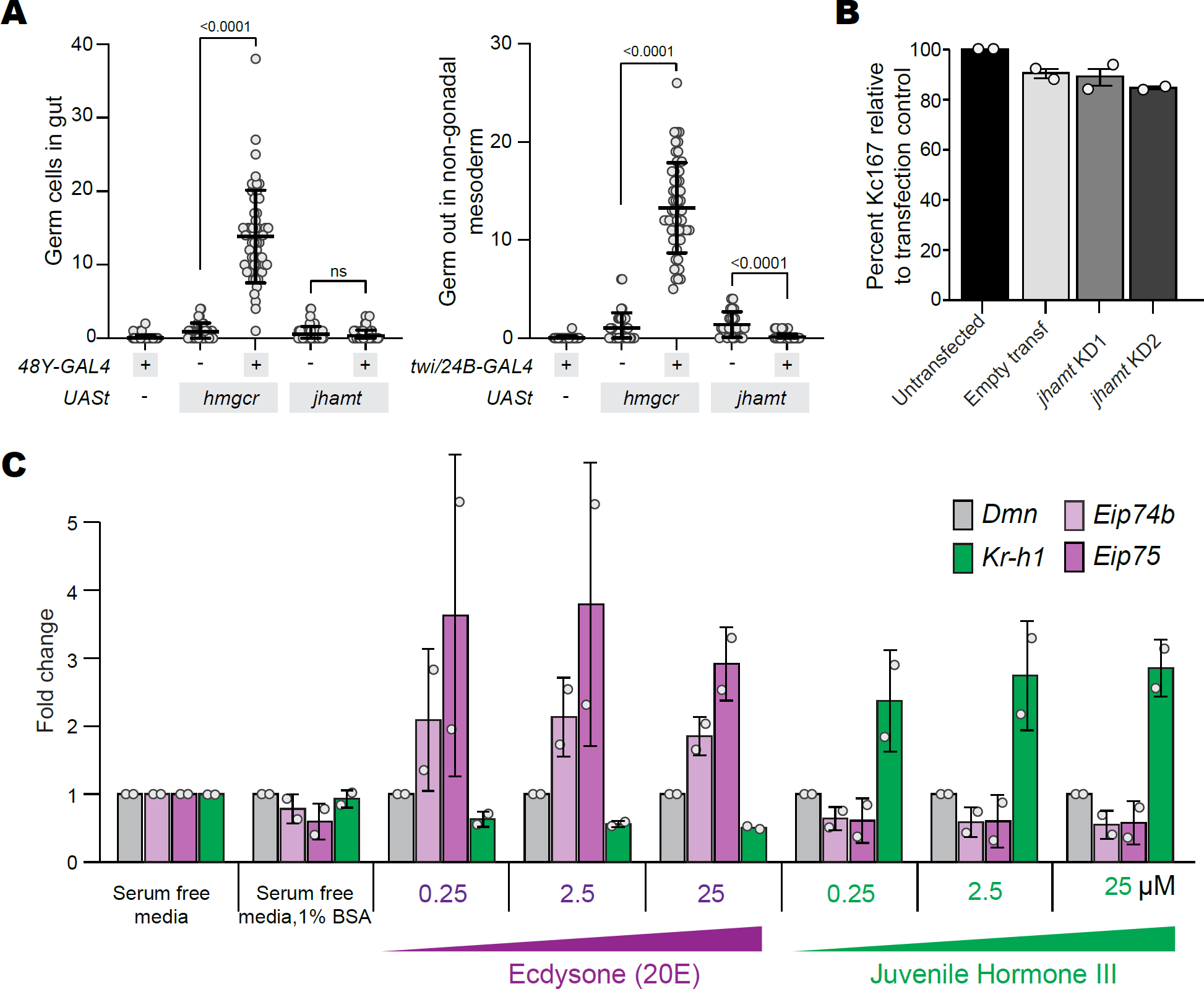
**A** *Left:* Quantification of the number of PGCs in Stage 14-16 embryos that remained in the gut upon ectopic expression of either *Hmgcr* or *jhamt* using the *48Y-GAL4* driver. *Right:* Quantification of the number of germ cells in Stage 14-16 embryos that contact Zfh-1-positive cells upon the expression of either *Hmgcr* or *jhamt* using the *twist(twi)-GAL4* and *24B-GAL4* drivers. Each dot represents one embryo. Bars and whiskers represent median +/-IQR and statistical significance was analyzed by the Mann-Whitney U test. **B** Quantification of percent Kc167 cells, used to condition serum-free media, upon *jhamt* mRNA knocked down relative to transfection control. Bars and error bars represent mean +/-standard deviation from two biological replicates, each including at least two technical replicates. **C** Quantification of the fold change in mRNAs in S2 cells upon incubation with varying concentrations of 20-Ecdysone (20-E) or JH III for six hours prior to mRNA isolation. mRNAs from two verified ecdysone-responsive genes were measured: *Eip74b* (lighter purple) and *Eip75* (darker purple) and one JH-responsive gene was measured, *Kr-h1* (green) and normalized to the housekeeping gene, *DCTN2-p50*. Error bars represent standard deviation from two biological replicates, each consisting of at least two technical replicates.

**Supplemental Data for Figure 4.**
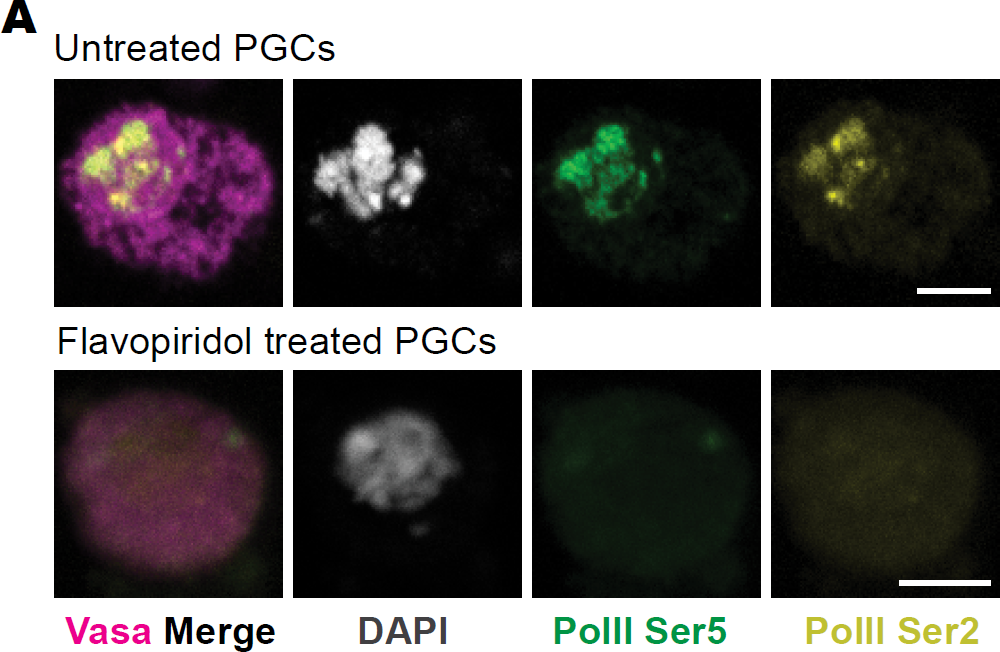
Confocal images of *Drosophila* PGCs enriched by FACS and stained for Vasa (magenta), RNA Pol II, Ser5p (green), RNA Pol II, Ser2p (yellow) and DAPI (gray). Bottom panel shows a germ cell treated with 1µM Flavopiridol for 3 hours. Scale bars represent 5µm.

**Supplemental Data for Figure 5.**
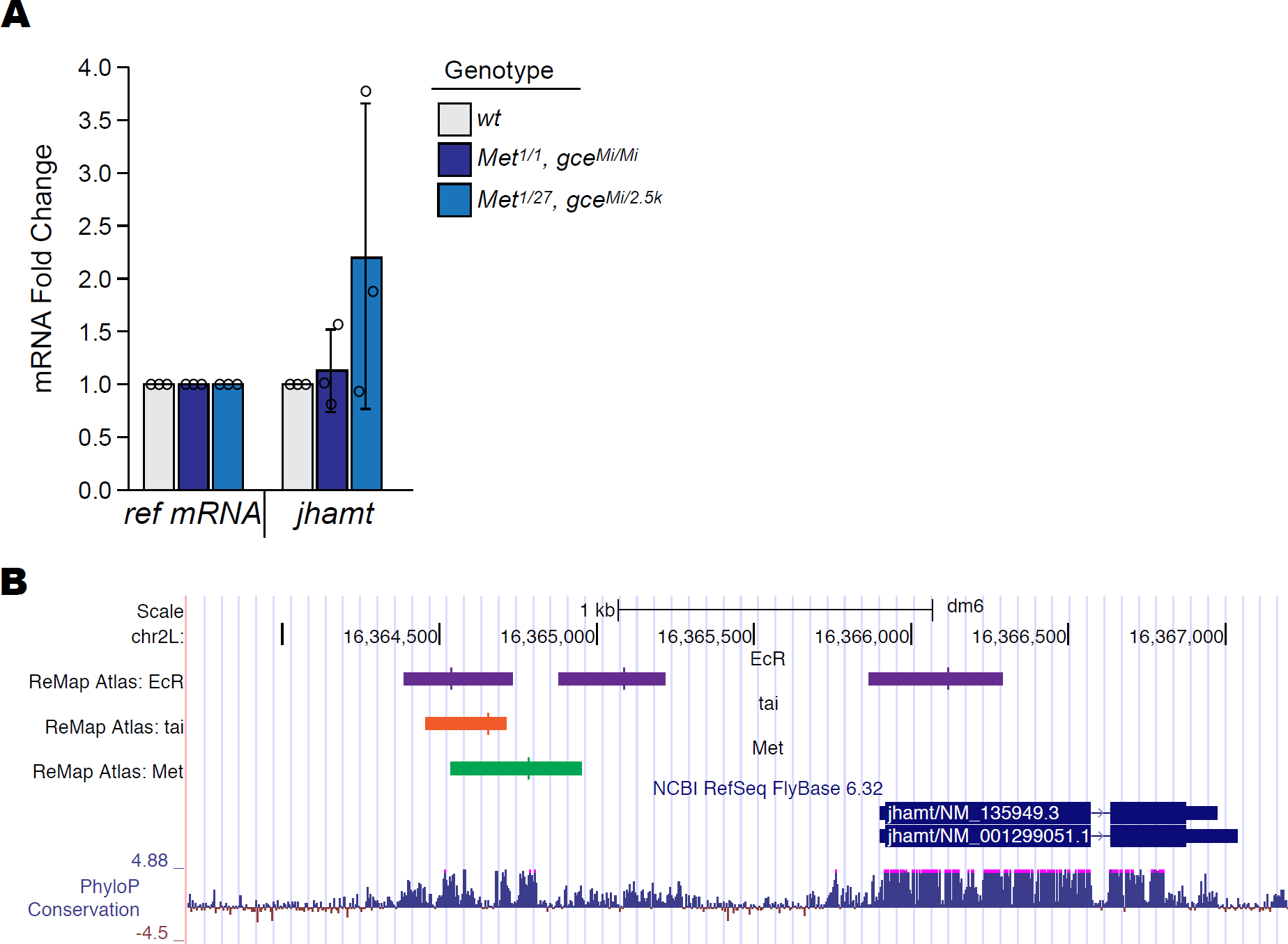
**A** Fold change of *jhamt* mRNAs in *Met, gce* mutant one-day old adult females relative to wildtype control animals. deltaCt was calculated relative to the average Ct of *DCTN5-p25* and *Und*. Shown are Fold Change of *jhamt* mRNA in adults homozygous for the newly recombined mutant alleles *Met*^1^*, gce^Mi^* (dark blue) and heteroallelic combination of *Met*^1^*, gce^Mi^*with existing or *Met*^27^*, gce*^2^.^5k^ alleles (light blue). Bars and error bars represent mean +/-standard deviation from three biological replicates, each containing pooled mRNA from five females. **B** Schematic of the Ecdysone Receptor (EcR), Taiman (Tai) and Met binding sites near the *jhamt* locus. Binding sites were obtained from ModEncode^63^.

**Supplemental Data for Figure 6.**
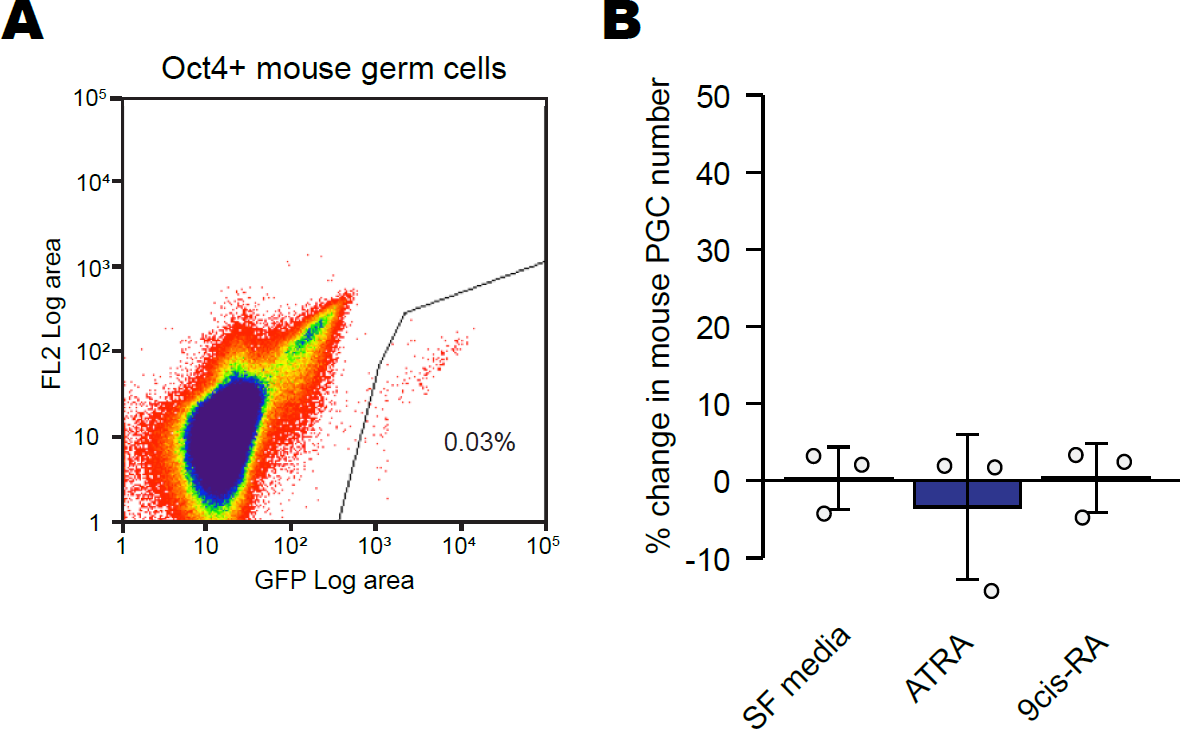
**A** Scatter plots from FACS of GFP-positive PGCs enriched from trunks of E10.5 mouse embryos carrying the *Oct4!1PE:GFP* transgene. The number inside the gate lines denotes the percentage of collected events in a typical biological replicate. **B** Quantification of the percent change in the number of mouse germ cells before and two hours after application of 1µm ATRA or 9cis-RA. Two hours was chosen because this is the duration of the *in vitro* migration assay. Graphed is the mean and standard deviation from three biological replicated. Each dot represents the averaged technical replicates of each biological replicate.

